# High resolution diel transcriptomes of autotetraploid potato reveal expression and sequence conservation among rhythmic genes

**DOI:** 10.1101/2025.04.22.650056

**Authors:** Ann Feke, Brieanne Vaillancourt, Kaitlyn Acheson, Julia Brose, John P. Hamilton, Dionne Martin, Yi-Wen Wang, Joshua C. Wood, C. Robin Buell, Eva M. Farre

## Abstract

Photoperiodic changes in diel cycles of gene expression are pervasive in plants. Timing of circadian regulators together with light signals regulate multiple photoperiod dependent responses such as growth, flowering or tuber formation. However, for most genes the importance of cyclic mRNA levels is less clear. We analyzed the diel transcriptome of modern cultivated potato, a highly heterozygous autotetraploid. Clonal propagation and limited meiosis have led to the accumulation of deleterious alleles and therefore tetraploid potato is an ideal model system to investigate the conservation of cyclic expression and cyclic genes during the artificial selection process. We observed that cyclic alleles were more highly expressed than non-cyclic ones and were highly co-expressed not only under diel cycles but also across tissues, developmental stages and stress conditions. Moreover, the smaller ratio of non-synonymous to synonymous differences within cyclic as compared to non-cyclic allelic groups indicates that cyclic genes, in general, have more conserved core functions than those of non-cyclic ones. In accordance with this observation, fully rhythmic allelic groups were highly enriched in photosynthesis and ribosome biogenesis genes, which play core functions in plants. Furthermore, we investigated differences in cyclic expression patterns between photoperiods. We identified transcription factors potentially regulating the strong differences in phase between photoperiods observed in ribosome biogenesis and pathogen response genes. Finally, analyses of genes involved in tuber formation suggests that the regulation of *CO* gene transcription is not the only factor enabling tuberization under long days in modern cultivated potato. This study not only provides high quality diel transcriptomic datasets of cultivated potato but also important insight on the role of allelic diversity in rhythmic expression in plants.

## BACKGROUND

Cultivated potato, *Solanum tuberosum* L. Group Tuberosum (2*n*=4*x*=48), is a heterozygous autotetraploid that is vegetatively propagated. Polyploidy has been associated with domestication and there is evidence that polyploidy preceded domestication for many crops [1]. In addition, variation in clock associated genes have also been linked to crop domestication [2–5] indicating that cyclic gene expression is an important factor targeted in this process.

Most of the of previous work on cyclic transcriptomics in polyploid crops has relied on haploid genome assemblies or on allopolyploids where variability can be attributed to ancestry such as *Brassica rapa* and hexaploid wheat. *B. rapa* is a mesohexaploid, generated through a whole genome triplication after it split from *Arabidopsis* 13-43 million years ago and a subsequent diploidization event [6–8]. Hexaploid wheat (AABBDD) formed through interspecific hybridization between a domesticated diploid and a diploid species 8,500-9,000 years ago [9]. In these cases, homeologs were able to evolve either before polyploidization in hexaploid wheat or after in *B. rapa*. In *B. rapa*, 42% of paralog pairs have differential expression patterns under light/dark conditions, and most display differences in median level of expression [10]. In hexaploid wheat, 64.15% of triads also display differences in rhythmicity patterns [11]. These results indicate functional divergence of these cyclic genes, which might be partly caused by their different origins. There is no information on how rhythmic expression correlates across alleles in autopolyploids such as potato.

Potato species have undergone more recent polyploidization events. Sexual polyploidization via 2n gamete production has been recurrent in wild as well as cultivated potato species [12]. Recent studies indicate that modern potato cultivation was initiated with tetraploid Andean landraces and that the breeding process has included admixing with tetraploid Chilean genotypes and wild species [5, 13]. Due to its poor fertility, caused by different factors including polyploidy [12], cultivated potato is primarily maintained vegetatively leading to an accumulation of deleterious alleles and inbreeding depression, resulting in a high genetic load [14]. Therefore tetraploid potato is a good model system to investigate the conservation of cyclic expression and cyclic genes during the artificial selection process.

Modern cultivated potatoes are able to grow in a wide range of latitudes [15] indicating flexibility to photoperiod adjustment for both physiological as well as developmental processes. In potato, CYCLING DOF FACTOR 1 (CDF1) is a repressor of *CONSTANS* (*CO*) genes, which inhibit tuberization under long days by activating the expression of *SP5G* (*SELF PRUNING 5G*) a repressor of *SP6A*, a phloem mobile signal that induces tuber formation [16, 17]. FLAVIN-BINDING KELCH REPEAT F-BOX 1 (FKF1) is an F-box protein that induces the degradation of CDF1 under short day conditions by interacting with the c-terminus of CDF1 [16, 18]. In modern cultivars, truncated *CDF1* alleles, encoding proteins that no longer interact with FKF1, accumulate and repress *CO* expression under both short and long day conditions, thus allowing for tuberization even under long photoperiods [5, 15]. It is believed that the circadian clock is also involved in this process by generating oscillations of expression of *FKF1*, *CDF1* and *CO* in an analogous manner to the regulation of flowering time in *Arabidopsis*. This current model of photoperiod control of tuberization was developed in *S. tuberosum* Group Andigena, which requires short day conditions to tuberize. It is not clear how photoperiod affects the expression of these genes in cultivated potato, which are able to tuberize under long days yet still retain some photoperiod sensitivity.

Our previous studies on cultivated potato showed that both leaves and tubers are able to maintain robust diel and free running rhythms [19]. However, these experiments were performed using unphased assemblies of the highly heterozygous tetraploid potato using 3’ Tag-seq, and thus lack the ability to distinguish the expression of the individual alleles. Here we performed a detailed analysis of diel expression under long and short photoperiods on the cultivated potato *S. tuberosum* Atlantic using a haplotype resolved genome assembly and annotation.

## METHODS

### Plant growth and sample collection conditions for diel datasets

*Solanum tuberosum* cv. "Atlantic" (referred to herein at "Atlantic") tissue culture plantlets were transplanted to soil (Suremix, Michigan Grower Products, Galesburg) in 14 cm deep x 9 cm square pots in and fertilized with Peter’s 20-20-20 weekly. Plants were grown in a BioChambers High-Light Hi/Lo CO2 GRC-40 growth chamber under either short days (SD), 12 h light (22°C, 350 μmol s^-1^ m^-2^), 12 h dark (18°C), or long days (LD), 16 h light (22°C, 350 μmol s^-1^ m^-2^), 8 h dark (18°C). Sampling occurred every 2 h for 24 h. Dawn samples (ZT 0, ZT 24) were collected in the dark and dusk samples (ZT 12) in the light. At each time point we collected tissue from three plants (replicates). For leaf tissue we collected three terminal leaflets from the third and fourth newest fully expanded leaves per replicate. Under short days we also collected tissue from the three largest tubers per plant, using a 5 mm punch and discarding epidermal tissue. All tissue was snap-frozen in liquid nitrogen, then stored at –80°C prior to RNA isolation. At the time of harvest, short day plants were 9 weeks old and long day plants 4.5 weeks old.

### Identification of syntenic allelic groups

Syntelogs were identified between the genomes of DM v6.1, Atlantic v3, *Solanum chacoense* v5, *Solanum candolleanum* v1, and *Solanum lycopersicum* M82 v1 using GENESPACE (v1.2.3) [65] with the representative working model proteins for each genome downloaded from SpudDB [65] and setting the ploidy to 1, 4, 1, 1, 1 respectively. The syntelog sets were extracted from the pangenome database file from each GENESPACE run.

Further processing was performed using a custom python script to select allelic groups with four or fewer alleles, requiring that alleles were found on the same chromosome but without duplication on a single haplotype. For genes with fewer than four alleles in phased chromosomes, we selected alleles from unphased chromosomes when available.

### Total RNA isolation, library preparation, and sequencing of diel expression datasets

Total RNA was isolated using a modified hot borate method [56] followed by treatment with TURBO DNase (Invitrogen, Waltham, MA) to remove any residual DNA. RNA integrity was verified via gel electrophoresis and Fragment Analyzer (Agilent Technologies, Santa Clara, CA). Short day RNA-Seq libraries were prepared using the Illumina Stranded mRNA Prep, Ligation kit (San Diego, CA) with IDT for Illumina Unique Dual indexes (Coralville, IA) and sequenced on the NovaSeq 6000 in paired end mode 100 nt by the Research Technology Support Facility Genomics Core at Michigan State University. RNA-Seq libraries for long day samples were prepared using the PerkinElmer NEXTFLEX Rapid Directional RNA-Seq Kit 2.0 with NEXTFLEX Poly(A) Beads 2.0 and NEXTFLEX® RNA-Seq 2.0 Unique Dual Index Barcodes (Waltham, MA) and sequenced on the NovaSeq 6000 in paired end mode 100 nt or 150 nt by the Texas A&M AgriLife Research: Genomics and Bioinformatics Service.

### RNA-Seq data processing of diel expression datasets

To check for contamination in sequencing data Kraken2 v1.2 [57] was run using the pre-built database k2_pluspfp_20220908 (https://benlangmead.github.io/aws-indexes/k2). RNA-Seq reads were then cleaned using Cutadapt v4.1 [58] to discard poor quality bases and adapter sequences by providing 3’ adapter sequences for trimming from both read one and two (-a, -A) along with the following parameters: -m 70 -q 30 --trim-n --times 2 -l 100. FastQC (https://www.bioinformatics.babraham.ac.uk/projects/fastqc/) and MultiQC [59] were both used to visualize quality and additional sequencing metrics for all libraries before and after cleaning. Kraken2 results, percent of reads discarded by Cutadapt cleaning, percent GC content, and over- represented sequence(s) outliers were flagged and individual libraries discarded based on results. After Cutadapt cleaning if a library had greater than 40 million read pairs then reads were subsampled using seqtk v1.3-r106 (https://github.com/lh3/seqtk) with the sample function and the parameters -s100 40000000. Transcript abundances were quantified using Kallisto quant v0.48.0 [60] with the Atlantic v3 high confidence representative gene models [20], the --rf-stranded option, and a k-mer size of 21. Pearson correlation coefficients were calculated between biological replicates using R “pairwise.complete.obs”, method “pearson”.

### Atlantic Developmental Gene Expression Atlas

A replicated developmental and stress gene expression atlas of the *S. tuberosum* Atlantic spanning 16 tissues and 8 treatments (https://spuddb.uga.edu/expression.shtml) were obtained from NCBI under BioProject PRJNA753086). Reads were evaluated for quality using FastQC(https://www.bioinformatics.babraham.ac.uk/projects/fastqc/) and MultiQC [59]. Reads were cleaned using Cutadapt (v4.2, v4.4) [58] with a 3’ adapter provided for both read one and read two, a minimum read length (-m) of 40 nt, low quality reads trimmed from the 3’ end of each read (-q 30), flanking N’s removed (--trim-n), and the number of times to trim an adapter set to two (--times 2). Reads were checked for contamination using Kraken2 (v2.1.) [57] with the database k2_pluspfp_20220908. Subsampling was then performed using seqtk (v1.3/1.4; https://github.com/lh3/seqtk) based on the median total reads after cleaning for each species, ranging from 43 to 80 million reads. To determine expression abundances, cleaned subsampled reads were used with Kallisto quant (v0.4.8) [60] and their respective set of high confidence representative gene models with the --rf-stranded option and the k-mer size set to 21.

### Normalization of RNA-seq experiments

RNA-seq read counts were normalized using the rlog function from the R package DEseq2 [61]. We defined expressed transcripts as those with at least one sample with an rlog > 0. All three photoperiodic datasets (short day leaf, short day tuber, and long day leaf) were co-normalized to allow for comparisons in expression level across samples. Tissue samples and stress samples from the Developmental Gene Expression Atlas were normalized separately. The samples, Closed Flower, Cold Leaf Control, Hooked Stolon, Immature Fruit, Mature Fruit, Open Flower, Root Control, Sprout, Stem Control, Swollen Stolon, Tuber S1, Tuber S2, Tuber S3, Tuber S4, Tuber S5 and Young Leaf 10am, were included in the ’Tissue’ dataset and samples Methyl Jasmonate Control, Methyl Jasmonate Treatment, BTH Control, BTH Treatment, Salt Leaf Control, Salt Leaf Treatment, Cold Leaf Control, Cold Leaf Treatment, Drought Leaf Control, Drought Leaf Treatment, Heat Control, Heat Treatment, Root Control, Salt Root Treatment and Drought Root Treatment included in the ’Stress’ dataset. The diurnal time course samples from the Developmental Gene Expression Atlas were not included in this study. Normalized gene expression can be found in Dataset S2-4.

### Rhythmicity analysis

For the analysis of rhythmicity, rlog normalized expression values were analyzed using the JTK-CYCLE algorithm in the MetaCycle r-package [62] using the parameters of minper=20, max_per=26, and adjustPhase="predictedPer". Transcripts were considered rhythmic if the JTK *p*-value was less than 0.001. Phase of rhythmic gene expression were used as provided by JTK when phase < 24, and circularized (phase_circularized_ = phase_JTK_ – 24) when phase ≥ 24. Rhythmicity data be found in Dataset S5-7.

### Determination of correlation of gene expression

Normalized gene expression profiles generated as described in “Rhythmicity Analysis” were further normalized using *z-*scoring using the python package scipy (1.11.3) [63] to compare for the daily patterns of expression rather than the abundances. *z*-score normalization was performed within each individual time course or experiment, rather than across experiments. The Pearson correlation of the *z*-scored expression patterns was calculated for each pair of alleles using the series.corr() method in the python package pandas (2.1.1) [64]. When classifying allelic pairs by rhythmicity for the correlations based on the tissue and stress datasets from the Atlantic Developmental Gene Expression Atlas, the rhythmicity values from the LD datasets were used since these experiments were performed under 15 h light/9 h dark conditions. Correlation data be found in Dataset S8-10.

### Differential Expression analysis

Differential expression between photoperiods was performed using the R-package DEseq2 [61]. For this analysis, the time component of RNAseq samples was disregarded, and samples were assigned to the photoperiod from which they originated. Transcripts were considered differentially expressed if the Benjamini-Hochberg false discovery rate adjusted *p-*value was < 0.05 and the abs(log2Fold Change) >1. Differential expression data be found in Dataset S11-12.

### Tissue specificity analysis

We calculated the Tau index as described by [22]. We defined expressed transcripts as those whose average rlog across replicates was larger than zero in at least one tissue. We set tissues with rlog equal or smaller than zero as not expressed, i.e. equal to zero.

### Determination of synonymous and nonsynonymous substitution rates

We used SynMap within the CoGe online software suit (https://genomevolution.org/coge/) using default settings to calculate synonymous (Ks) and nonsynonymous (Kn) substitution rates between syntenic alleles in Atlantic [67]. We performed syntenic analysis using SynMap using the annotation file ATL_v3.working_models.gff3 available in spuddb.uga.edu [66]. We then filtered the syntenic allelic pairs determined using our previous syntenic analysis. Only Ks values larger than 0.001 and smaller than 3 were used for the Kn/Ks ratio calculations.

### Functional enrichment

We first determined the functional protein annotation using Mercator4 v7.0 [68] (www.plabipd.de/mercator_main.html) and the Atlantic high confidence representative gene models (SpudDB) (Dataset S13). Functional enrichment was done using Mercator4 BIN enrichment analysis online tool (https://www.plabipd.de/mercator_main). The selection of background genes for the enrichment analyses depended on specific research question and is provided in the text and/or the respective figure legend. Over-representation was calculated using a one sided Fischer’s exact test. The FDR-adjusted *p*-value cutoff was set to 0.05.

### Promoter cis-element enrichment analysis

Identification of binding sites within Atlantic promoters was performed using the annotatePeaks.pl program from the HOMER suite [69]. In brief, candidate binding sites from equivalent transcription factors from *Arabidopsis thaliana* were downloaded from the Plant Transcription Factor Database [70] and converted into the HOMER motif format, prior to being compared to the promoter regions of the Atlantic genome. Promoter regions were defined as 1,500 bases upstream to 100 bases downstream of the annotated transcription start sites. Fisher’s Exact Tests were performed using the python package scipy [63] in order to determine whether promoters of query genes were enriched for these binding sites.

### Identification of circadian clock, photoperiod and tuberization associated syntenic groups

Using our computational pipeline, we associated 460 of those CPT alleles to 138 allelic groups. We determined that the main reason for the lack of allelic group identification for 39 of our CPT transcripts was copy number variation and therefore, manually associated them to ’putative syntenic allelic groups’ (Table S2). For several of these genes (*CONSTANS*, *EARLY FLOWERING 4*, *SELF PRUNING 5G LIKE*, *SUGARS WILL EVENTUALLY BE EXPORTED TRANSPORTER*), syntenic allelic groups were likely not identified using our automatic pipeline as these genes had local tandem duplications in the Atlantic genome. In addition, the allelic groups of *CO*s, *ELF4* and *SPA1* (*SUPPRESOR OF PHYA-105*) contain currently annotated non-syntenic duplicates (Table S2). For example, in addition to the syntenic triplet of *CO* genes on haplotype 2 of chromosome 2 there is an additional ’triplet’ of *CO* genes, which we named *CO1b*, *CO2b, CO3b*. To investigate whether these additional haplotype specific copies were annotated in other cultivated potato genomes, we searched for non-syntenic orthologs in the genomes of Otava [23] and Cooperation-88 [24] using BLASTP. We did not find any non-syntenic copies of CPT genes in these genomes. These results indicate that these additional copies in Atlantic may be mislocalized in the current Atlantic haplotype assembly or represent cultivar-specific structural variation. After this manual curation we linked the 499 CPT gene models to 154 allelic groups in Atlantic, of which, 53% had four alleles, 25% had three and only 14% and 8% had two and one allele respectively.

### Graphs and statistical analysis

Graphs were generated in R version 4.3.1, with the exception of cis-element motifs. If not otherwise indicated, statistical tests were implemented in R version 4.3.1 and described in the respective figure legends.

## RESULTS

### Fully rhythmic allelic groups display stronger rhythms and higher gene expression in both leaves and tubers

We used synteny between in *S. tuberosum* cv. “Atlantic” (referred to herein as “Atlantic”) haplotypes and the genomic reference *S. tuberosum* Group Phureja DM 1-3 516 R44 (DM) to identify high confidence syntenic allelic groups or genes (Dataset S1). Of these genes, 2,827 (11.9 %) had a single allele in the our Atlantic genome, 3,871 (16.3 %) had two alleles, 5,436 (22.9 %) had three alleles, and 11,599 (48.9 %) had four alleles, similar to the previous reported distribution [20]. We defined ’expressed’ genes as those with at least one expressed allele.

To address the nature of differences in diel expression across haplotypes in cultivated tetraploid potato, we generated short day (SD, 12-hour light/12-hour dark) and long day (LD, 16-hour light/8-hour dark) 24 hour gene expression time courses in Atlantic during the tuber bulking phase. Leaf tissue was collected from soil-grown plants every 2 hours and used to construct RNA-Seq libraries in which the resulting reads were aligned to the phased assembly of the Atlantic genome [20]. Tuber tissue was collected only under the short day photoperiod.

Differences in the phase of expression or overall expression level between alleles could have masked cycling expression patterns in the non-haploid resolved genome of Atlantic [20]. Using the haploid resolved Atlantic genome assembly, we first identified rhythmic transcripts using the JTK algorithm [21] (Datasets S2-4). We selected one representative transcript model for each allele of each gene, and transcripts were considered rhythmic if they resulted in an adjusted *p*- value of less than 0.001 (Fig. 1A). We selected this stringent *p*-value to ensure high accuracy in the estimation of cyclic parameters, such as phase. Using these criteria we found 30,986 rhythmic in the leaf in either one or both of our conditions, corresponding to 33.6% of all expressed transcripts. Of these strong rhythmic transcripts, 8,591 (27.7 %) were identified in both SD and LD experiments. In contrast, in tubers, only 996, 1.1% of the expressed transcripts, cycled. Even among cyclic transcripts, those cycling in the tuber had larger adjusted *p*-values than those cycling in leaf tissue (Supplemental Fig. 1A). Most (889) of the tuber transcripts were also expressed in leaves and 52.3% of tuber rhythmic transcripts also cycled in leaf tissue in SD (Fig. 1A).

**Figure 1.**
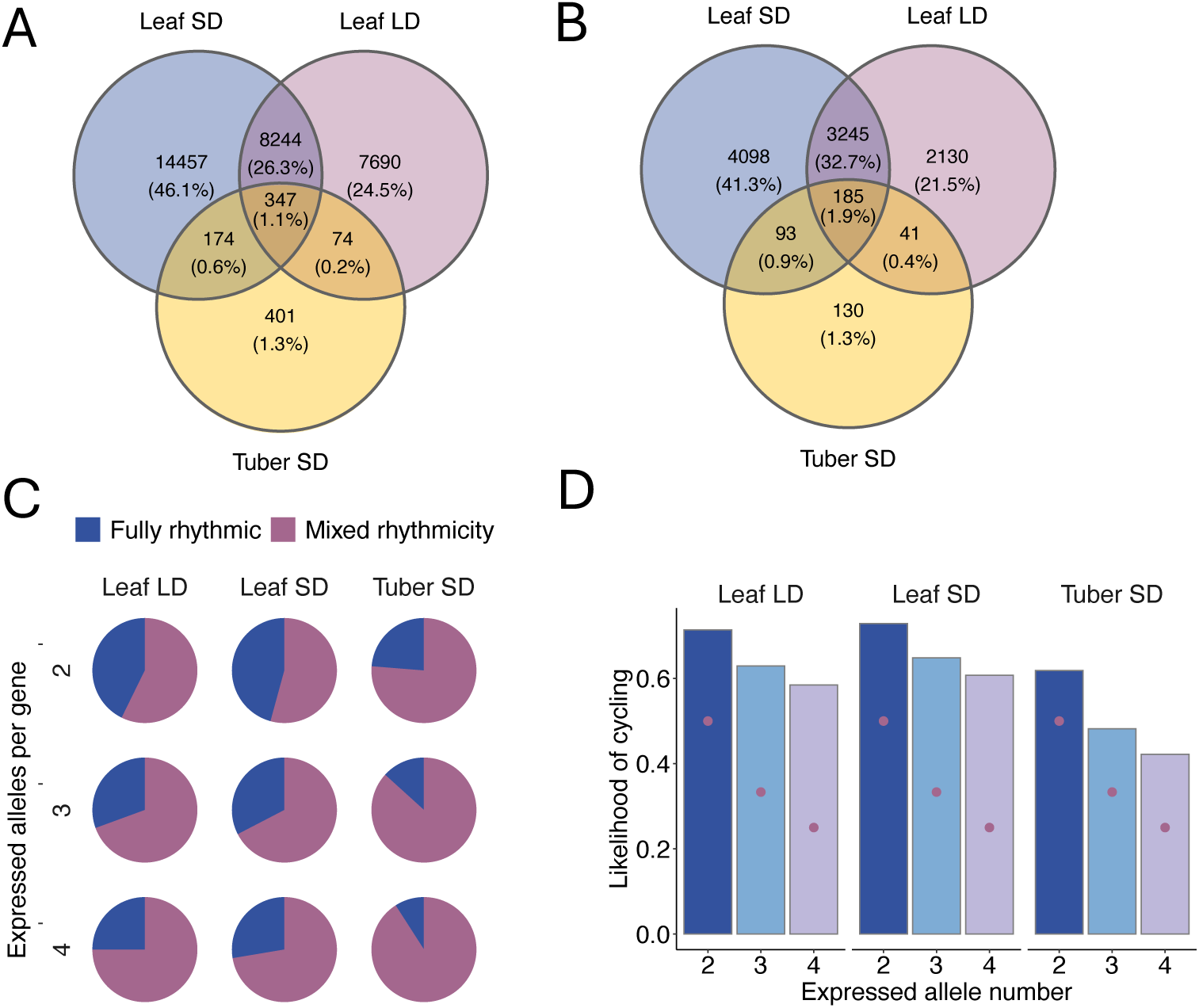
Diel cycling rates in cultivated potato. **A.** Number of cycling transcripts per condition. In parenthesis, the percent cycling rate. **B.** Number of cycling genes per condition. In parenthesis, the percent cycling rate. **C.** Distribution of cycling genes with either all expressed alleles cycling (Full rhythmicity) or at least one not cycling allele (Mixed rhythmicity). **D.** Likelihood of one allele cycling within cyclic genes with 2-4 expressed alleles. Dark purple dots represent minimum likelihood of cycling in the respective category, e.g. for a cyclic gene with two expressed alleles the minimum likelihood of cycling is 0.5.

We defined cyclic genes, as those with at least one cyclic allele. Overall, gene rhythmic rate across different conditions was similar to the rate of rhythmicity among transcripts (Fig. 1A, B). Structural variation among the four haplotypes occurs [20] and we tested whether cyclic genes had a different number of alleles than non-cyclic ones. There was only a very minor difference in the number of alleles of cyclic and non-cyclic genes, with an average 3.5 expressed alleles for non-cyclic and 3.3 for cyclic genes (*p*-value < 2.2e-16, Kruskal-Wallis rank sum test).

We next asked how many genes exhibited "mixed rhythmicity", i.e. they have at least one non-cyclic and one cyclic allele. We observed that the majority of rhythmic genes with at least two expressed alleles existed in a mixed rhythmicity state, rather than being fully rhythmic (all expressed alleles were rhythmic) (Fig. 1C). Genes with more alleles were more likely to have mixed rhythmicity in both leaves and tubers (Fig. 1C). In tubers, there was a slightly higher percentage of genes with mixed rhythmicity, which might be due to the lower robustness of tuber cyclic expression (Supplemental Fig. 1A). These observations agree with previous studies in tetraploid potato cultivars showing that genes with more alleles have higher variation in expression [20]. However, the likelihood of a transcript being rhythmic was still higher if it belonged to a cyclic gene, such that cyclic genes contained more cyclic alleles than the expected minimum (Fig. 1D). These results indicate that rhythmicity might be relatively well conserved between alleles.

Alleles from fully rhythmic genes had more robust rhythmicity than rhythmic alleles of genes with mixed rhythmicity, as quantified using the adjusted *p*-value of our JTK analyses (Fig. 2A). This observation suggests that the more cyclic alleles a gene contains the more important rhythmic expression might be for its function. Alleles in fully rhythmic genes (all four alleles rhythmic), when compared to all cyclic alleles in multiallelic genes, were enriched in functions related to the light reactions of photosynthesis and ribosome biogenesis (Fig. 2B, Supplemental Table S1), indicating that rhythmic mRNA levels might be especially important for these functions.

**Figure 2.**
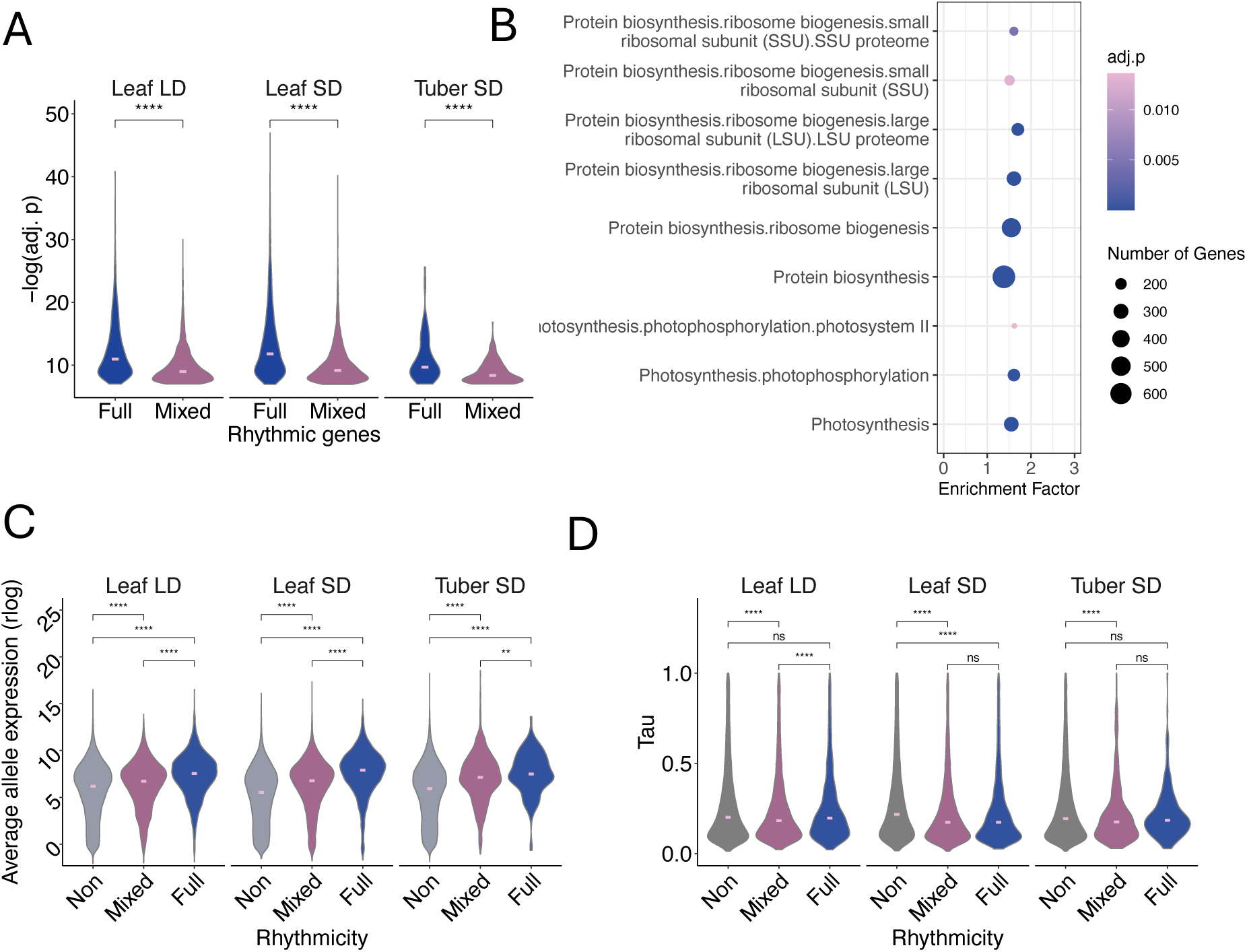
Allele specific cycling in potato tissues. **A.** Cycling strength, as determined by JTK calculated FDR adjusted mean p-value of cyclic transcripts in genes with either full or mixed rhythmicity. SD, short days; LD, long days. **B.** Functional enrichment of transcripts within fully cyclic genes in comparison to all cyclic transcripts. **C.** Average allele expression within genes with no cyclic alleles (No), at least one cyclic allele (Mixed) or all alleles cycling (Full). **D.** Degree of tissue specific expression as determined by Tau within genes with no cyclic alleles (No), at least one cyclic allele (Mixed) or all alleles cycling (Full). For A,C and D: Wilcoxon signed-rank test with Bonferroni correction was used for the statistical analyses, such that adjusted p-value < 0.0001 (****), <0.001 (***), < 0.01(**), <0.05 (*) and ns (not significant); horizontal bar indicates average. For C and D Kruskal-Wallis multiple groups test *p*-value < 0.0001.

Finally, diel rhythmicity was also associated with higher expression. Transcripts from cyclic genes were higher expressed than those from non-cyclic genes (Fig. 2C). Additionally, within mixed-rhythmicity allelic pairs, the cyclic allele was, on average, more highly expressed than the non-cyclic one (Supplemental Fig. 1B). It is possible that low expression is associated with higher noise, reducing the sensitivity of rhythmicity detection algorithms specifically for lower expressed transcripts. Alternatively, as it has been hypothesized in tetraploid wheat, that arrhythmic homeologs are silenced and therefore are not rhythmic [11]. Regardless, we still observed significantly higher expression of cyclic alleles than non-cyclic ones amongst well-expressed transcripts (average rlog > 5, Supplemental Fig. 1C).

We then tested whether cyclic transcripts might also be more widely expressed across tissues and developmental stages non-cyclic transcripts. We used a publicly available allele-specific dataset of Atlantic, composed of samples from 16 different tissues and developmental stages, including stolon, tuber root, stem, leaf, flower, fruit and sprout tissues to calculate the Tau index [22]. A higher Tau index indicates stronger tissue specific expression. Non-rhythmic genes and non-rhythmic alleles in allelic pairs had slightly more tissue specific expression than non-rhythmic ones (Supplemental Fig. 1D, Fig. 2D). For example, ∼8.5% of non-rhythmic transcripts in leaf tissue had a Tau larger than 0.8, which indicates a high tissue specificity, but only ∼4.5% of rhythmic ones did.

### Rhythmic alleles display high correlation of expression across multiple tissues and stress conditions

We investigated whether there were differences in the timing of expression between rhythmic alleles. Most fully rhythmic allelic pairs (>81%) displayed phase differences of less than 2 h, which was our sampling time resolution (Fig. 3A). Only 6-7% of those pairs had phase differences greater than two hours. To further assess the similarity of timing of expression rhythms within genes we calculated the correlation of expression between allelic pairs using z-score normalized expression. We observed strong correlation of expression among pairs with two rhythmic alleles (Fig. 3B), such that more than 80% of those pairs had a correlation of expression greater than 0.85. In contrast, allelic pairs with either none or only one rhythmic allele had significantly weaker correlations. For example, only 28-34% of pairs with only one rhythmic allele had correlations larger than 0.85. Taken together, we observed that although the majority of rhythmic allelic groups have mixed rhythmicity, the alleles that cycle display a high degree of rhythmic coherence. Thus, our data suggests that, despite the significant sequence and structural changes that occur within potato homologous chromosomes and alleles [20, 23–25], the timing of expression of rhythmic genes is highly conserved across alleles.

**Figure 3.**
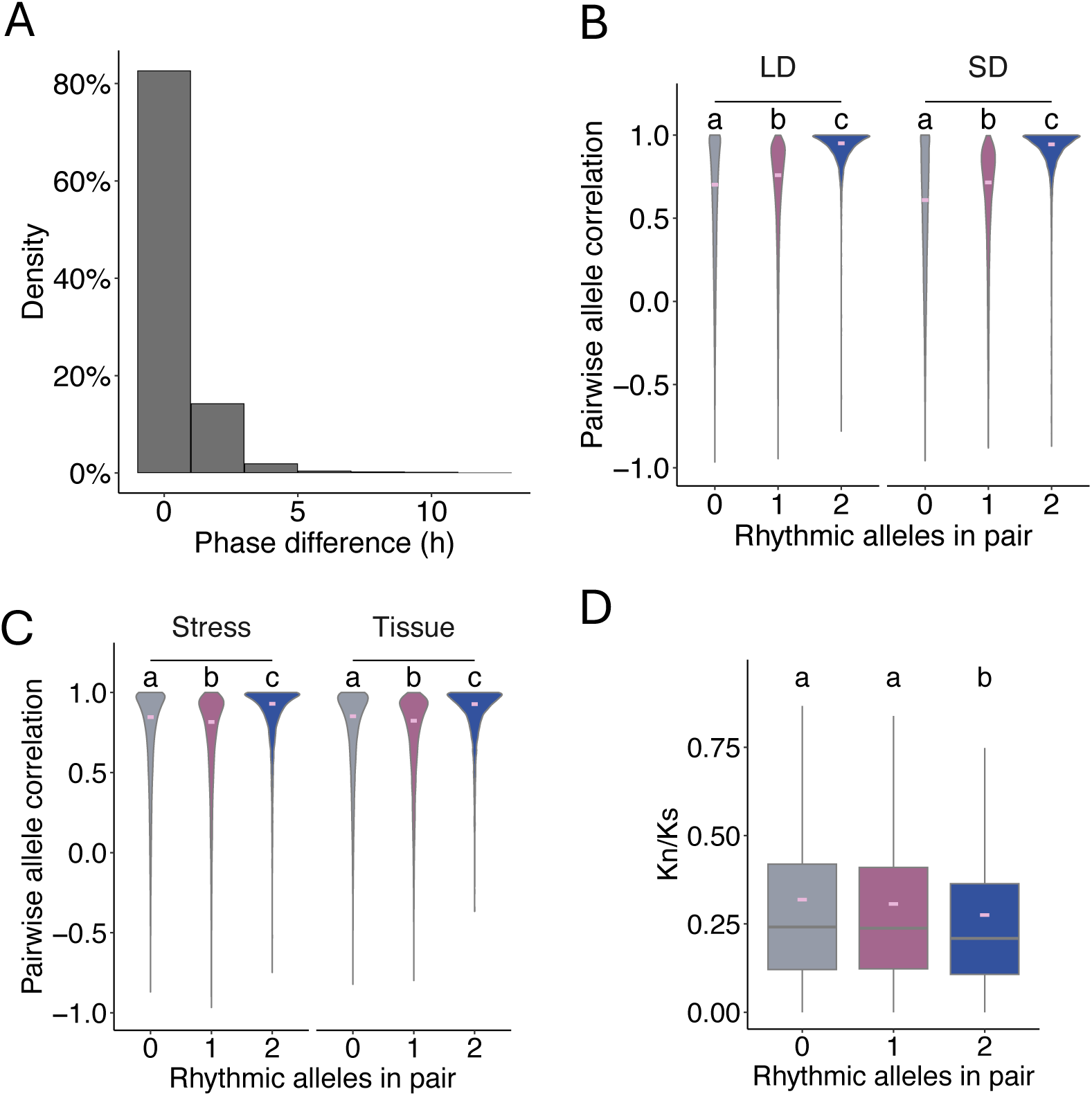
Similarities of expression among cyclic alleles. **A.** Phase differences between cyclic alleles. Data contains all three datasets, leaf (short day and long day) and tuber (short day). **B.** Correlation of z-score normalized gene expression between allelic pairs in leaf tissue. SD, short day; LD, long day. Rhythmicity was determined in the respective (SD or LD dataset). **C.** Correlation of z-score normalized gene expression between allelic pairs across different tissues (Tissue), or stress/hormone treatments (Stress). Rhythmicity was determined in the LD dataset, since the the Tissue and Stress experiments were performed under 15 h light photoperiods. **D.** The ratio of non-synonymous (Kn) to synonymous (Ks) substitution rates between allelic pairs. Rhythmicity was determined in the SD dataset. **B-D,** letters indicate significant differences adjusted *p*-value < 0.0001, of post-hoc Wilcoxon signed-rank test with Bonferroni correction. For all three plots Kruskal-Wallis multiple groups test *p*-value < 0.0001.

To investigate the tight co-expression of rhythmic allelic pairs within single tissues corresponds to a wider degree of co-expression across tissues and stresses, we used the tissue and developmental expression dataset used for our tissue specificity analysis (’Tissue’ dataset), and another publicly available dataset composed of eight hormone and abiotic stress treatments and their respective controls (’Stress’ dataset). Interestingly, co-expression of fully rhythmic allelic pairs was higher for all datasets than for pairs in which in which one or fewer alleles cycled (Fig. 3C). For example, in these tissue and stress datasets ∼70% of fully rhythmic pairs had correlations larger than 0.85, but only ∼ 45% of mixed rhythmicity pairs did.

To estimate whether these highly coexpressed cyclic allelic pairs are under purifying selection, we examined the ratio of nonsynonymous (Kn) to synomymous (Ks) substitution rate within our syntenic allelic pairs. The distribution of Ks between these pairs in Atlantic was similar to that of other tetraploid potatoes and independent of their rhythmicity [25](Supplemental Fig. 2). We observed a smaller Kn/Ks ratio within fully rhythmic syntenic allelic pairs, than within pairs with either none or only one rhythmic allele (Fig. 3D). This observation indicates that fully cyclic allelic pairs might maintain higher functional similarity in addition to high expression similarity than non-cyclic genes.

### Tuber phase is delayed with respect to the leaf under temperature cycles

Our high temporal resolution data sets enabled us to perform accurate phase determinations between leaves and tubers under short day conditions. Transcript expression peaked at all times throughout the day (Fig. 4 A-C), although in leaves most rhythmic transcripts peaked at dawn or during the second half of the light period, with fewer transcripts peaking in the late night (ZT18- 22). In the tuber, the majority of cyclic genes peaked during the light period. Interestingly, of the 521 transcripts rhythmic in both leaves and tubers, 47% displayed at least a 2 h delay (Fig. 4D), with an average delay of 4.4 h. In contrast, only 19% of those rhythmic transcripts had an advanced phase equal or larger than 2 h in tubers with respect to leaves. Functional categories enriched among transcripts with delayed phase in tuber included circadian, photoperiod and light signaling, and proteasomal degradation (Supplemental Fig. 3). Differences in phase between above ground and below ground tissues have been reported in *Arabidopsis thaliana* [26–29].

**Figure 4.**
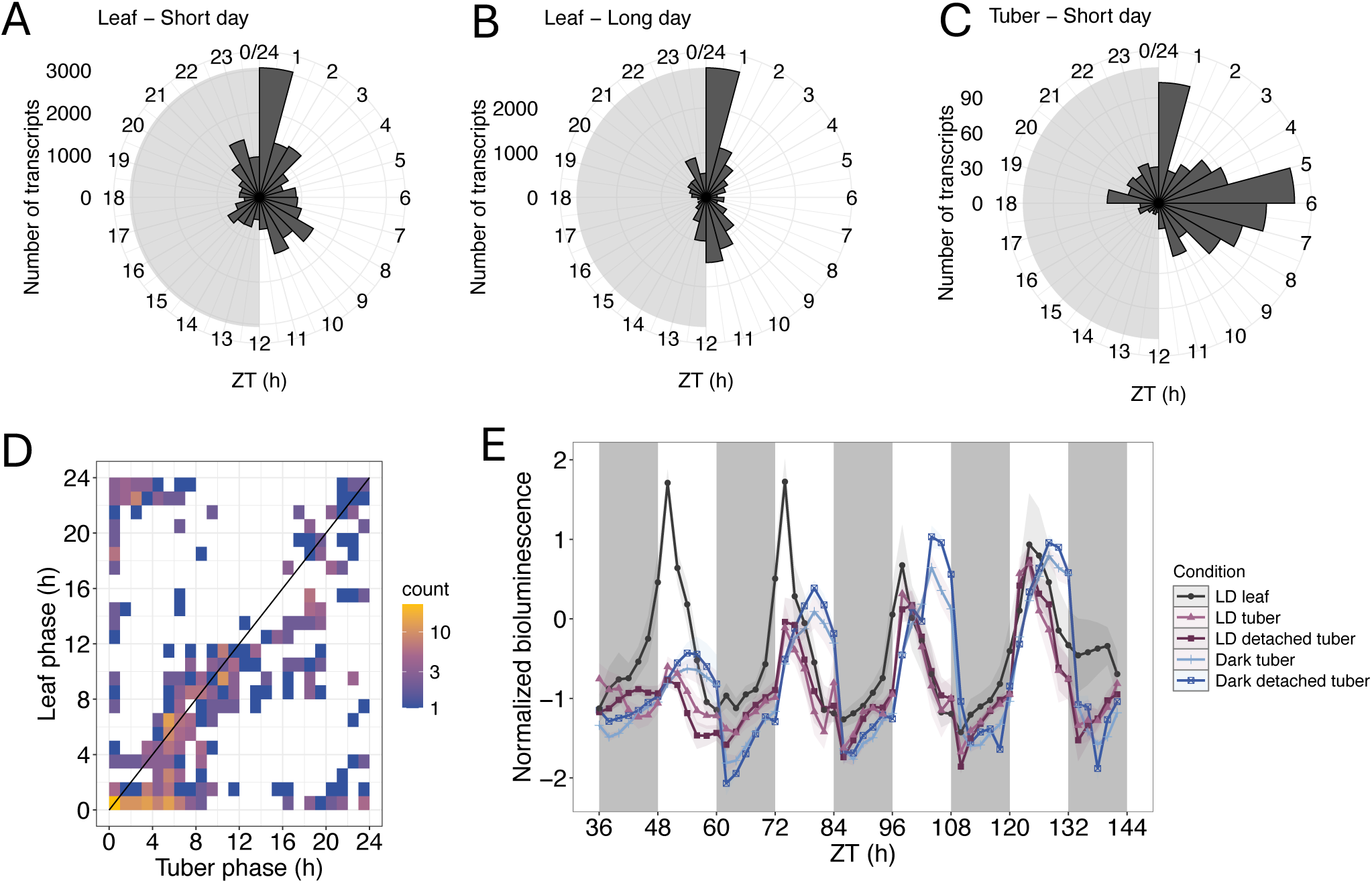
Changes in timing of expression across tissues and photoperiods. **A.** Distribution of phases of gene expression under short day conditions in leaves. **B.** Distribution of phases of gene expression under long day conditions in leaves. **C.** Distribution of phases of gene expression under short day conditions in tubers. **D.** Comparison of phase of expression of genes cycling in both leaves and tubers under short day conditions. **E.** Expression of the *GH3::LUC* bioluminescence reporter (average ± SE, n = 6-16). Grey areas indicate dark/subjective dark periods. LD, light/dark conditions.

Since our plants were grown under both light and temperature cycles, phase differences between underground tubers with respect to leaves could be due to delays of the entrainment signals coming from the shoot, delays in the perception of environmental temperature oscillations due to the temperature buffering capacity of the soil, or differences in entrainment phase in the soil due to the absence of direct light signals. We used the previously characterized expression reporter, *GH3::LUC,* to determine the phase of expression in leaves and tubers [19]. We observed that there were no significant differences in phase between leaves, and attached or detached tubers when both tissues were exposed to the same light and temperature cycles (Fig. 4E). However, when tubers were kept in the dark, the phase of both attached and detached tubers was delayed with respect to leaves, even if both tissues were exposed to temperature cycles. These results indicate that light signals are able to entrain the tuber clock directly and in their absence there is a delay in phase in that tissue with respect to leaves. Under natural field conditions temperature cycles are offset from light/dark cycles such that the minimum temperature generally coincides with dawn and the maximum temperature with the end of the day. This shift, together with temperature buffering by the soil, would suggest that tuber rhythms of field grown plants might be several hours phase shifted from leaf rhythms.

### Photoperiod dependent changes in gene expression in potato leaves

We next evaluated photoperiod dependent changes of rhythmic gene expression in potato leaves. Among transcripts rhythmic in SD and LD, we observed that a higher percentage had significantly elevated average expression under short days (∼11%) than under long days (∼5%) (Fig. 5A). We also observed that the average amplitude of gene expression was larger in our SD dataset than under LD (Fig. 5B). However, only ∼23% of transcripts with larger amplitudes in SD had significant elevated expression under short photoperiods (Supplemental Fig. 4A). The larger amplitude could explain why we detected a higher number of rhythmic transcripts in SD (Fig. 1A,B). Transcripts with larger amplitudes under short days were expressed almost exclusively in during the light in both photoperiods (Supplemental Fig. 4B) and, in comparison to other cyclic transcripts, were strongly enriched in photosynthesis-related functions (FDR-adjusted *p*-value < 0.00001) (Supplemental Table S1). Among the transcripts encoding for transcription regulators with larger SD amplitudes we identified transcripts encoding for REVEILLE (RVE), CYCLING DOF FACTOR (CDF), HEAT SHOCK TRANSCTRIPTION FACTOR A6B (HSFA6B) and TCP proteins (Supplemental Fig. 4C).

**Figure 5.**
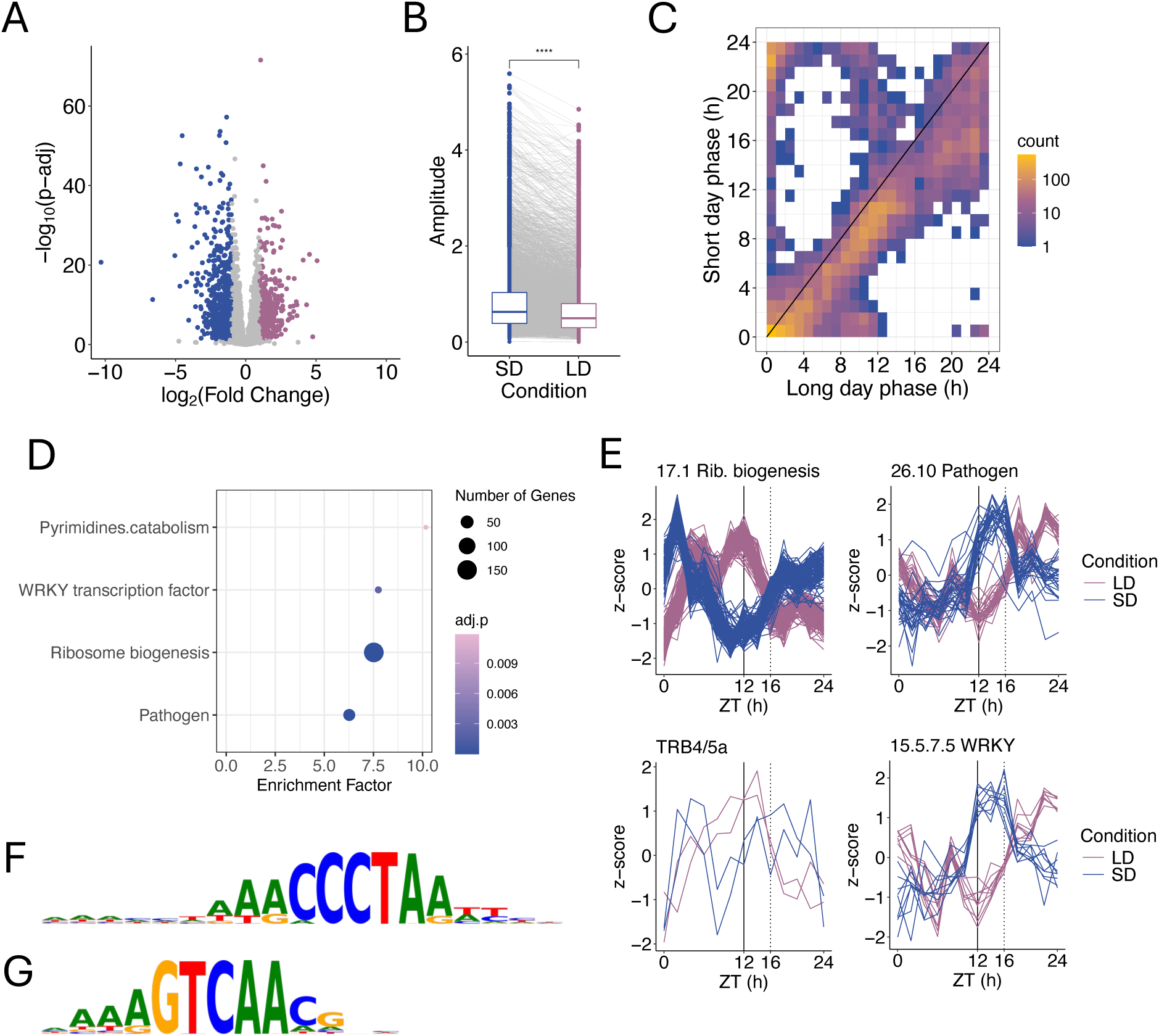
Expression changes between short (SD) and long photoperiods (LD). **A.** Volcano plot of differentially expressed transcripts between photoperiods. Blue indicates upregulated expression under SD and pink under LD. **B.** Pairwise amplitude (determined by JTK) of transcripts rhythmic in SD and LD. (****) Indicates adjusted *p*-value < 0.0001, Wilcoxon signed-rank test with Bonferroni correction. **C.** Phase of expression in short and long days among genes cycling in both conditions in potato leaves. **D.** Functional enrichment of genes with changes in expression of 6 h under long day with respect to short day in potato leaves, when compared to all genes rhythmic in both SD and LD. **E.** Expression pattern of genes in the MapMan4 17.1 Ribosome biogenesis category with large changes in phase. Note that two TRB4/5a transcripts (Soltu.Atl_v3.03_0G008090 and Soltu.Atl_v3.03_0G008150) have the same sequence and therefore the same estimated expression. TRB4/5a-like transcript, Soltu.Atl_v3.03_0G008210 was not plotted, because it was very low expressed. Expression pattern of genes in the MapMan 26.10 Pathogen and the 15.5.7.5 WRKY transcription factor categories with large changes in phase. Expression values were z-score normalized before plotting, the average of three biological replicates is shown. All times represent hours after dawn (ZT). **F.** Summary motif for TRB-binding sites enriched in the Ribosome biogenesis category of panel C. **G.** Summary motif for WRKY-binding sites enriched in genes in the Pathogen category of panel C.

As in other studies [30] we observed that many rhythmic genes had delayed phases under long days relative to short days (Fig. 5C). About 51% of genes that cycled in both photoperiods were delayed in phase by at least 2 h under longer days, with an average delay of 3.7 h. We did not find any significant differences in functional enrichment in those transcripts when compared to all transcripts rhythmic in both conditions. In contrast, only about 12 % of cycling genes had an advanced phase under long days. Strong differences in phase in expression between photoperiods has been hypothesized to be a mechanism to optimize translation under lower energetic resources under short days in photosynthetic organisms [31]. We found 716 genes with phase delays under long days of at least 6 h. These genes were strongly enriched in functions related to ribosome biogenesis, pathogen responses, pyrimidine catabolism and WRKY transcription factor activity (Fig. 5D, Supplemental Table S1).

Protein biosynthesis genes counted for 28% of the 716 transcript with at least 6 h phase delay under long days, most of which were related to ribosome biosynthesis (Supplemental Table S1). The mRNA level of these genes peaked around dawn under short days and the second half of the light period under long days (Fig. 5E). Moreover, although not all were detected as exhibiting phase shifts, in our experiments, the majority of transcripts in the functional categories ribosome biogenesis-large ribosomal subunit 17.1.2 and ribosome biogenesis-small ribosomal subunit 17.1.3 followed similar expression patterns (Supplemental Fig. 5). Similar photoperiod dependent changes in expression have been observed in some ribosomal genes in *Arabidopsis thaliana* [32]. In *Arabidopsis,* TELOMERE BINDING PROTEINS (TRB) have been shown to bind to the promoters of ribosome biogenesis genes via telobox motifs [33]. We identified several TRB genes in potato (Supplemental Fig. 6), one of which, TRB4/5a, displayed maximum mRNA levels at the end of the day in long days and around dawn under short days (Fig. 5E). We also observed that TRB-related *cis*-elements were enriched in upstream regions of the ribosomal genes with strong changes in phase (Fig. 5F, Supplemental Table S1). Taken together, these photoperiod controlled TRB proteins could be involved in the observed photoperiod dependent shift in expression.

Of the 76 genes related to pathogen responses that were rhythmic under both photoperiods, 41 displayed at least 6 h delays in phase under long days. These genes included 20 genes potentially involved in effector-triggered immunity and 10 genes related to systemic acquired resistance (Supplemental Table S1). These pathogen response genes had peaks of expression at the beginning of the night period under short days and end of the night under long days (Fig. 5E).

Interestingly, we found that WRKY transcription factors were also enriched within strong phase shifted genes and had a very similar expression pattern as the pathogen related genes. Moreover, *cis*-element motifs recognized by WRKY transcription factors in *Arabidopsis* were enriched in the upstream regions of the phase shifted pathogen related genes (Fig. 5G), indicating that the WRKY genes displaying changes in phase are involved in pathogen responses in potato.

### Expression characteristics of clock, photoperiod and tuberization associated genes

Diel and circadian regulation of gene expression plays a key role in the regulation of growth and photoperiod responses in plants, and are hypothesized to be involved in tuber formation in potato [16, 18, 34]. We identified 499 potential circadian, photoperiod or tuberization (CPT) transcripts in the Atlantic genome based on similarity to *Arabidopsis thaliana* genes (Supplemental Table S2). Using our computational pipeline and manual curation we linked the 499 CPT gene models to 154 allelic groups in Atlantic, of which, 53% had four alleles, 25% had three and only 14% and 8% had two and one allele respectively.

We classified our CPT genes into five categories based on their characterized function in *Arabidopsis*, potato or tomato. The ’Clock’ set of genes included genes associated with core clock function such as the pseudoreponse regulators (*PRR*s), or *EARLY FLOWERING TIME 3*. ’Clock_Aux’, included genes with pleiotropic functions that can affect circadian control, such as casein kinases or phytochrome interacting factors. The ’Clock_photoperiod’ category includes genes involved in both clock and photoperiod control such as *GIGANTEA* (*GI*). The ’Photoperiod’ category includes genes mainly involved in photoperiod signaling such *CO* or *CDF*s. Finally, the ’Tuberization’ category includes genes involved in tuber formation and growth, such as *BEL*-like transcription factors, gibberellin oxidases and genes involved in starch accumulation.

We observed that CPT genes in all categories displayed a higher percentage of rhythmicity and had higher rates of full rhythmicity than the genome wide averages (Fig. 6). Genes associated with core circadian clock function or photoperiodic regulation had a higher percentage of rhythmic transcripts than groups merely associated with the tuber formation process (Fig. 6A). Interestingly, the group of genes we named clock auxiliary had a lower number of rhythmic transcripts than the group of core clock genes in all conditions. Genes associated with core clock function were also more likely to have all their alleles rhythmic than genes in the other categories (Fig. 6B). Among the core clock associated allelic groups that were fully rhythmic in both leaf and tubers we found *REVEILLE 3/5* (RVE3/5), *RVE4/8*, *RVE1a*, *PRR5*, *PRR3a*, *PRR3b*, *LUX ARRHYTHMO/BROTHER OF LUX* (*LUX-BOAb*) and *LATE ELONGATED HYPOCOTYL* (*LHY*).

**Figure 6.**
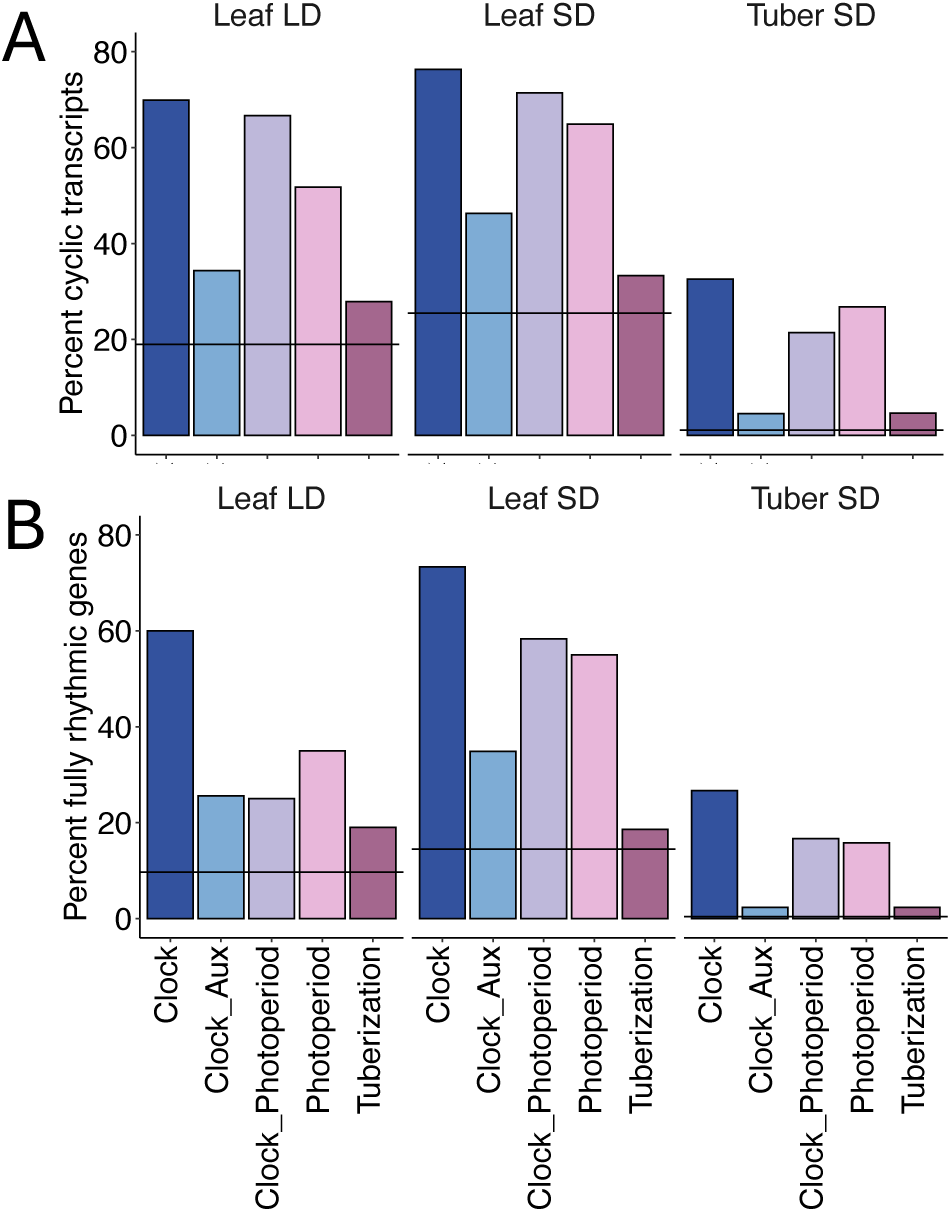
Rhythmicity of clock, photoperiod and tuberization related genes. Rhythmicity was quantified using JTK. Fully rhythmic genes were genes with all their alleles cycling. Horizontal lines indicate the respective genome wide percentages in each condition.

### Expression of photoperiod regulating genes in cultivated potato suggest the presence of a CO independent mechanism of SP6A induction

As other modern potato cultivars, Atlantic tuberizes under long photoperiods. In photoperiod sensitive potato (*S. tuberosum* Group Andigena), CO is a repressor of tuberization and its expression is repressed by CDF1 [16, 18]. The Atlantic genome contains four *CDF1* alleles. Two alleles (haplotype 1 and haplotype 0) are identical and encode for truncated proteins missing the C-terminus required for interaction with the F-BOX1 protein (FKF1) in the Arabidopsis ortholog (Supplemental Fig. 8). This interaction has been hypothesized to mediate CDF1 degradation under long days. Therefore, truncated CDF1 proteins are likely to have enhanced protein stability and are hypothesized to mediate the repression of *CO* transcription even under long days [18].

We observed that *CO1* and *CO2*, the highest expressed *CO* genes in Atlantic, displayed phase differences between photoperiods and slightly lower expression under short days in spite of the presence of deregulated CDF1 alleles (Fig. 7). This phase difference has also been observed in other potatoes, both photoperiod sensitive and less sensitive species [16, 35]. In contrast, the lower expressed *CO3* had slightly higher expression under long day conditions (Fig. 7).

**Figure 7.**
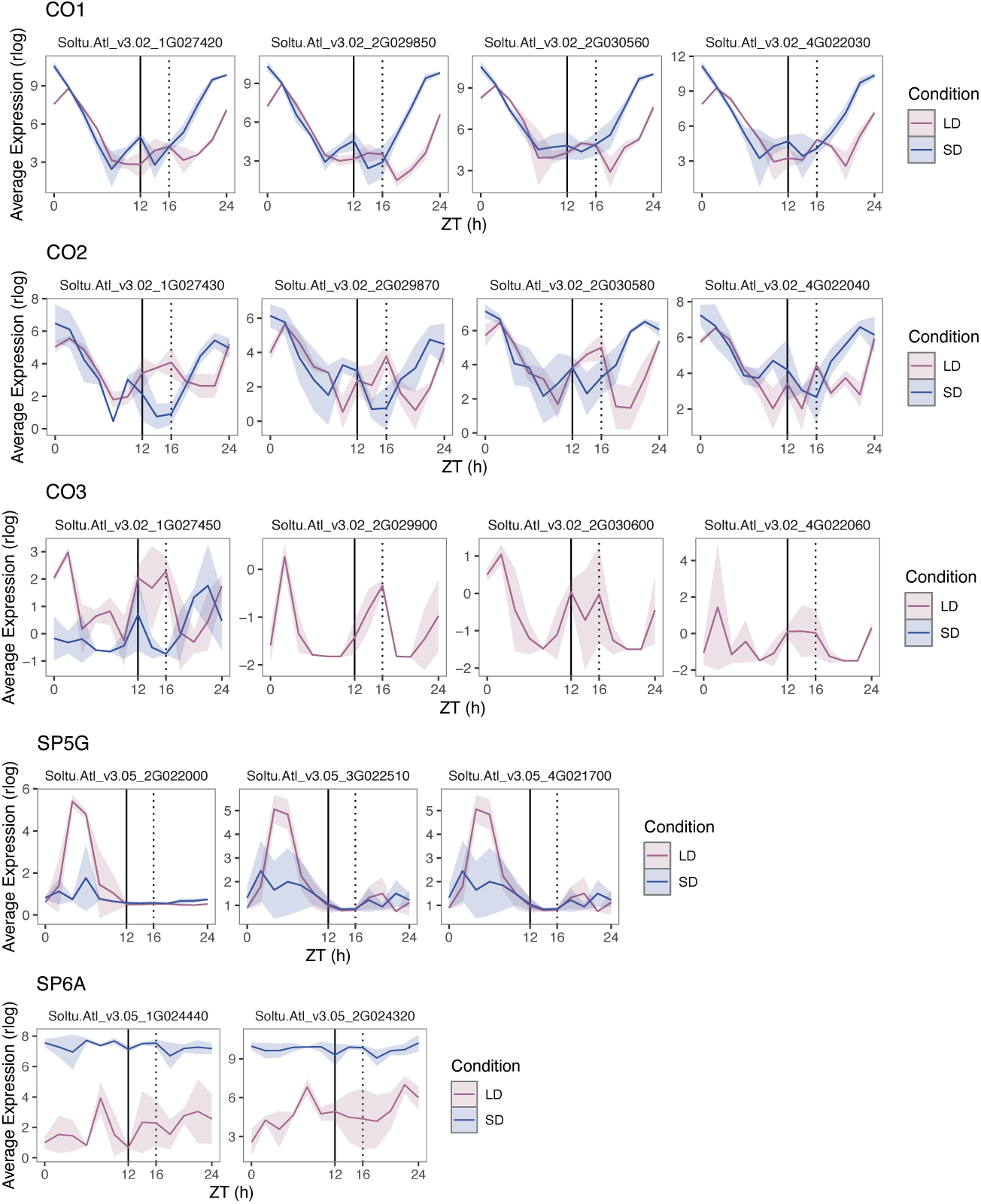
Expression patterns under short and long photoperiods. Data represents mean expression (average ± standard deviation, n = 3 ). SD, short days; LD, long days. *CO* genes include *CO1b* (Soltu.Atl_v3.02_2G030560), *CO2b* (Soltu.Atl_v3.02_2G030580), *CO3b* (Soltu.Atl_v3.02_2G030600).

The average expression of Atlantic *SP5G* and *SP6A* genes was significantly sensitive to photoperiod (Fig. 7). COs are believed to activate *SP5G* expression, which in turn represses *SP6A*, the mobile tuber inducing factor [16, 17]. There are four alleles of *SP5G* currently annotated in the Atlantic genome, three of which were expressed in the leaf, at a low level. All four alleles were predominantly expressed in tubers in our experiments (Supplemental Fig. 8). In leaves, *SP5G* alleles were more than four-fold higher expressed under long days. The two *SP6A* alleles found in Atlantic were about 30-fold higher expressed under short days than under long days. However, *SP6A* expression was still detectable in long days in Atlantic leaves, which might be sufficient to drive tuberization in this cultivar under those conditions.

## DISCUSSION

### Conserved highly expressed genes are more likely to display cyclic expression

Polyploidy in plants represents a good model to investigate relative short term selection on gene neofunctionalization and the importance of rhythmic expression for gene function. Here we observed high similarity of expression between alleles under diel cycles. Ferrari et al. [36] demonstrated that the timing of orthologous gene expression is strongly conserved across taxa. We had previously shown that cyclic gene expression to be predominantly retained after gene duplication [37]. We observed high conservation of cyclic expression within potato alleles, such that most cyclic allelic pairs displayed negligible differences in phase and high correlations gene expression under diel cycles (Fig. 3). These results indicate that the timing of gene expression is a key part of the function of cycling genes and is retained across different evolutionary scales.

Moreover, Ferrari et al. [36] had shown that genes conserved throughout the archaeplastida were more likely to cycle than expected by chance. We observe that rhythmic allelic pairs have smaller Kn/Ks ratios and therefore are more likely to have the same function than non-cyclic allelic pairs, indicating higher functional conservation. Potato allelic pairs display highly similar patterns of expression across tissues and multiple stress conditions than not cyclic allelic pairs (Fig. 3). Taken together these results support the hypothesis that cyclic gene expression occurs more often in conserved genes playing core roles and that there is less neofunctionalization of cyclic genes than non-cyclic genes.

We observed that rhythmicity was associated with higher expression across genes and also within alleles (Fig. 1C, Supplemental Fig. S1B). Similarly, cyclic paralogs in *B. rapa* [10] and cyclic homeologs in hexaploid wheat [11] are higher expressed than not cyclic ones. It has been shown that in yeast and in animals cyclic mRNA levels are associated with energetically ’expensive’ genes, i.e. genes with high costs of transcription and translation mainly because cyclic genes have higher expression [38]. Experiments in yeast, showed that cyclic expression amplitude increased in the presence of glucose, when cells had strong metabolic activity. The authors of these studies, therefore, hypothesized that cyclic expression enables the generation of high gene expression at time of need but still minimizing the overall cost of protein production [38].

Interestingly, in plants there might also be a correlation between stronger rhythms of higher amplitude with higher growth and metabolic rate. For example, in both gymnosperms and angiosperm trees diel oscillations only occur in the summer and not in the winter [39, 40]; young *Arabidopsis* plants have a greater number of cyclic genes and higher amplitude rhythms than in older plants [41], and pathogen responses that restrict growth are also associated with a decrease in amplitude [42]. Unicellular eukaryotic Archaeplastida also have higher amplitude rhythms of gene expression than multicellular ones, which may be linked to a higher growth rate [36]. In contrast, we observed a higher number of cyclic transcripts and an increase in amplitude under short photoperiods in cultivated potato (Fig. 1, Fig. 5). This increase in amplitude was mainly due to changes in photosynthesis related genes. In *Camelina sativa*, although biomass accumulation is decreased under short days, photosynthesis rates were increased, which has been hypothesized to compensate for the shortened light period [43]. Higher amplitude rhythms could enable this rate increase without additional cost of producing more RNA. Further studies are needed to test the role of metabolic activity on rhythmic gene expression in photosynthetic organisms.

### Some genes display specially strong changes in the phase of expression between photoperiods

It has been proposed that genes expressed at dawn display a decrease in their respective protein concentration as the photoperiod becomes longer due to the increases in vegetative growth rates in long days [31]. In contrast, evening expressed genes maintain or increase protein abundance with longer photoperiods. We observed that most ribosome biogenesis genes in cultivated potato displayed a strong difference in phase between photoperiods, such that mRNA content peaked around dawn under short days and at the end of the day under long days. In *Arabidopsis*, ribosome associated genes have similar photoperiod dependent changes in expression patterns [32] and appear to be preferentially translated at the end of the day under long photoperiods [44], indicating that, at least in long days, the phase of mRNA correlates with translation rate. The shifts in phase of expression of ribosomal genes could represent a mechanism to maintain protein content and therefore plant growth under long days in cultivated potato and other plants. The mechanism by which these changes in gene expression are regulated is yet unclear. As in *Arabidopsis*, we observe several *TRB* related genes that display similar changes in phase. *TRB* genes have been associated with ribosomal gene transcription, but further studies are needed to test whether they are also involved in the photoperiod changes observed [33, 45].

In our experiments, more than half of the transcripts in the functional category ’external stimuli response-pathogen responses’ that displayed cyclic expression had very strong differences in phase, such that their mRNA peaked early in the night under short days but at the end of the night under long days. Interestingly these genes were enriched in potential effector-triggered-immunity and systemic acquired resistance function. Light signals and the circadian clock have been shown to modulate time-dependent changes in sensitivity to pathogens in *Arabidopsis* [46–49]. In cultivated potato, the upstream sequences of these photoperiod regulated pathogen-response genes contained *cis*-elements that potentially mediate regulation via WRKY transcription factors. We identified three WRKY factors tightly co-expressed with these pathogen-related genes that are strong candidates for controlling these responses. They belong to three genes that previous studies in *S. tuberosum* Group Phureja named *StWRKY30*, *StWRKY38*, *StWRKY63* [50]. Potato WRKY transcription factors have been associated with stress responses including pathogen responses [50, 51]. However, there is currently no direct evidence for the involvement of these three WRKYs in pathogen responses in potato.

### Photoperiod control of tuberization in cultivated potato

Studies using different potato genotypes indicate that there are still unanswered questions on the CDF1-CO-SP5G-SP6A model of the photoperiod control of tuberization. Truncated alleles of *CDF1* are the principal mechanism for strongly reduced photoperiod sensitivity of tuberization in cultivated potatoes [5, 13, 18]. These truncated alleles lead to elevated levels of CDF1 protein and have been proposed to constitutively repress *CO* genes, leading to photoperiod independent tuberization [18]. Potato CO proteins, like *Arabidopsis* CO, are stabilized by light, and photoperiod-dependent differences in mRNA peak levels have been linked to large changes in CO protein between short and long days in photoperiod sensitive potato [16, 52, 53]. However, the characteristic phase difference of CO genes between short (peak at night) and long days (peak at dawn) is maintained in both photoperiod sensitive potatoes such as *S. tuberosum* spp Andigena and less photoperiod sensitive potatoes such as Atlantic or *S. neotuberosum* (Fig. 7) [35]. *S. neotuberosum*, a photoperiod insensitive potato, which contains truncated *CDF1* alleles, has very low levels of *SP5G* expression in both short and long photoperiods but also contain *SP6A* transcripts that are photoperiod responsive [35]. In contrast, our results show that Atlantic, which also contains truncated *CDF1* alleles, still displays photoperiod differences in both *SP5G* and *SP6A* expression (Fig. 7). Moreover, transgenic *S. tuberosum* spp Andigena plants overexpressing *CO1* have high levels of *SP5G* under both photoperiods but only display an increase in *SP6A* under short days and accordingly still display photoperiod sensitivity of tuberization [52]. These results highlight that we still do not have a full understanding of how the *CO* genes expression influences their respective protein abundances, and indicate that there might be a SP5G-independent mechanism of *SP6A* regulation in potato. BEL5, encoded by a phloem mobile mRNA, is also able to induce *SP6A* expression in potato leaves [54], however, in Atlantic leaves BEL5 expression is only induced about two-fold under short days, suggesting there is still an unknown layer of regulation of *SP6A* in short days (Supplemental Fig. S9). It has been proposed [18] that CDF1 could directly regulate *SP6A* in potato, however, *SP6A* has not been identified as a direct CDF1 target in recent work on cultivated potato [55]. Therefore, further studies are needed to understand how in potato CDF1 regulates tuberization in a *CO* independent manner.

## CONCLUSION

Our analysis of diel expression across haplotypes in cultivated tetraploid potato showed that rhythmic alleles not only retained high degree of co-expression under diel cycles but also across tissues, developmental stages and stress conditions, when compared to non-cyclic alleles. Moreover, smaller Kn/Ks ratios also suggest that rhythmic allelic pairs are likely to retain higher functional similarity. In general, rhythmic transcripts were higher and more constitutively expressed than non-rhythmic. Taken together these results suggest that rhythmic expression plays a key role on core plant cell functions. Finally, our results also showed that genes related to core circadian and photoperiod sensing functions were more likely to be rhythmic than genes involved in tuber formation. Moreover, photoperiod dependent changes of expression in genes associated with photoperiod sensing reveal open questions about the regulation of tuberization in cultivated potato.

## ACKNOWLEDGEMENTS

We thank Marisa Laitinen for help harvesting plants and tissue culture.

## DATA AVAILABILITY

The diel expression data can be accessed under the Sequence Read Archive BioProjects PRJNA957457 (short day) and PRJNA1093480 (long day). Atlantic Developmental Gene Expression Atlas expression data were obtained from NCBI under BioProject PRJNA753086.

## DECLARATIONS

### Ethics approval and consent to participate

Not applicable

### Consent for publication

Not applicable

## SUPPLEMENTAL TABLES

**Table S1.** Functional enrichment analysis determined using Mercator4. A. FullyRhymicGenes: Transcripts from multiallelic genes fully rhythmic in either SD or LD compared to all rhythmic transcripts from multiallelic genes in leaf tissue. B. TuberDelayed: Tuber cycling genes with delays of at least 2 hours with respect to all tuber expressed genes. C. LD_delayed_6h: Transcripts with phase delays of at least 6 hours with respect to short days compared to all transcripts rhythmic in both photoperiods. D. SDLD_Amplitude_Difference: Transcripts displaying increase amplitude under short than under long day conditions( > 0.5) with respect to all transcripts rhythmic under both photoperiods in leaves.

**Table S2.** Clock, tuberization, photoperiod genes (CPT). A. CPT_ATL_DM-syntenic associations between Atlantic and DM CPT genes. ’Gene’ indicates allelic group identified using the automatic pipeline (see Materials and Methods). ’Gene_manual’ contains additional associations after manual evaluation. H1-H4, indicate the haplotype, and H0 indicates gene models in unassembled scaffolds. B. CPT_C88_blast, five best blast hits of Atlantic CPT protein models against the Cooperation-88 (C88) protein models. C. CPT_Otava_blast, five best blast hits of Atlantic CPT protein models against the Otava protein models.

## LARGE DATASETS

**Dataset S1.** Allelic groups.

**Dataset S2**. Diel expression rlog

**Dataset S3**. Expression of Tissue samples from the Developmental Gene Expression Atlas

**Dataset S4**. Expression of Stress samples from the Developmental Gene Expression Atlas

**Dataset S5.** Leaf short day tissue cycling parameters as determined per JTK.

**Dataset S6.** Leaf long day cycling parameters as determined per JTK.

**Dataset S7.** Tuber short day cycling parameters as determined per JTK.

**Dataset S8.** Pairwise allelic expression correlations in short days.

**Dataset S9.** Pairwise allelic expression correlations in long days.

**Dataset S10.** Pairwise allelic expression correlations in Tissue samples from the Developmental Gene Expression Atlas.

**Dataset S11.** Pairwise allelic expression correlations in Stress samples from the Developmental Gene Expression Atlas.

**Dataset S12.** Differential expression short vs. long day determined by DEseq

**Dataset S13.** Differential expression leaf vs. tuber under short days determined by DEseq.

**Dataset S14.** Functional annotation of Atlantic using MapMan.

---

**Supplemental Figure 1.**
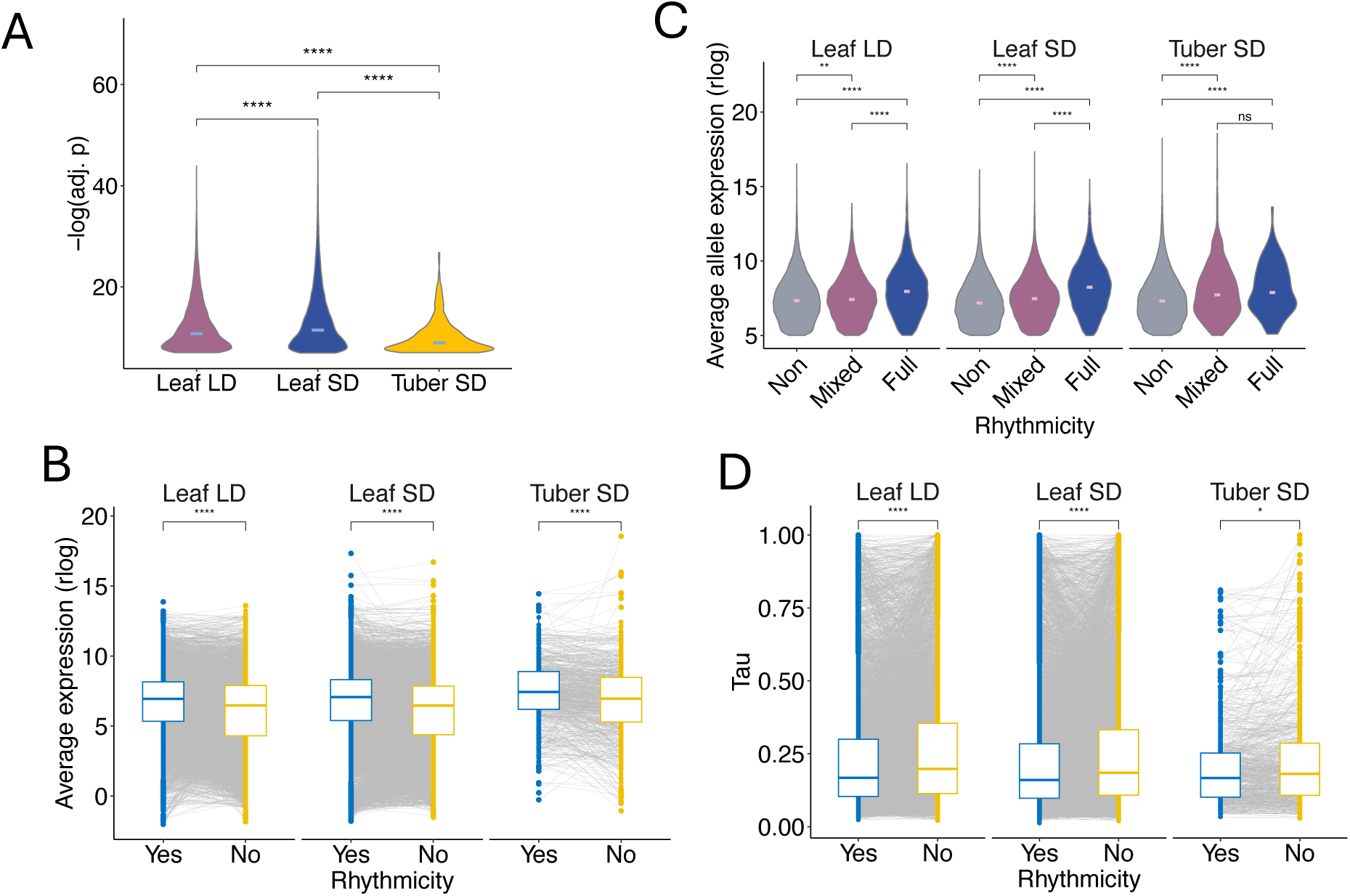
Allele specific cycling in potato tissues. A. Rhythmic strength, as determined by JTK calculated FDR adjusted p-value of cycling transcripts, in short day (SD) and long day (LD). Horizontal line indicates average. B. Comparison of average expression between allelic pairs, with one not cycling and one cycling allele. C. Average allele expression within genes with no cycling alleles (No), at least one cycling allele (Mixed), all alleles cycling (Full), of only highly expressed transcripts (rlog > 5). D. Tau index determining tissues specific expression of Atlantic transcripts either cycling or not cycling in the different conditions; Tau ranges between 0 (constitutively expressed) to 1 (tissue specific). Red bar indicates the mean. For A and C Kruskal-Wallis multiple groups test *p*-value < 0.0001. For A-D Wilcoxon signed- rank tests with Bonferroni correction were used for the statistical analyses, such that adjusted *p*- value < 0.0001 (****), <0.001 (***), < 0.01(**), <0.05 (*) and ns (not significant).

**Supplemental Figure 2.**
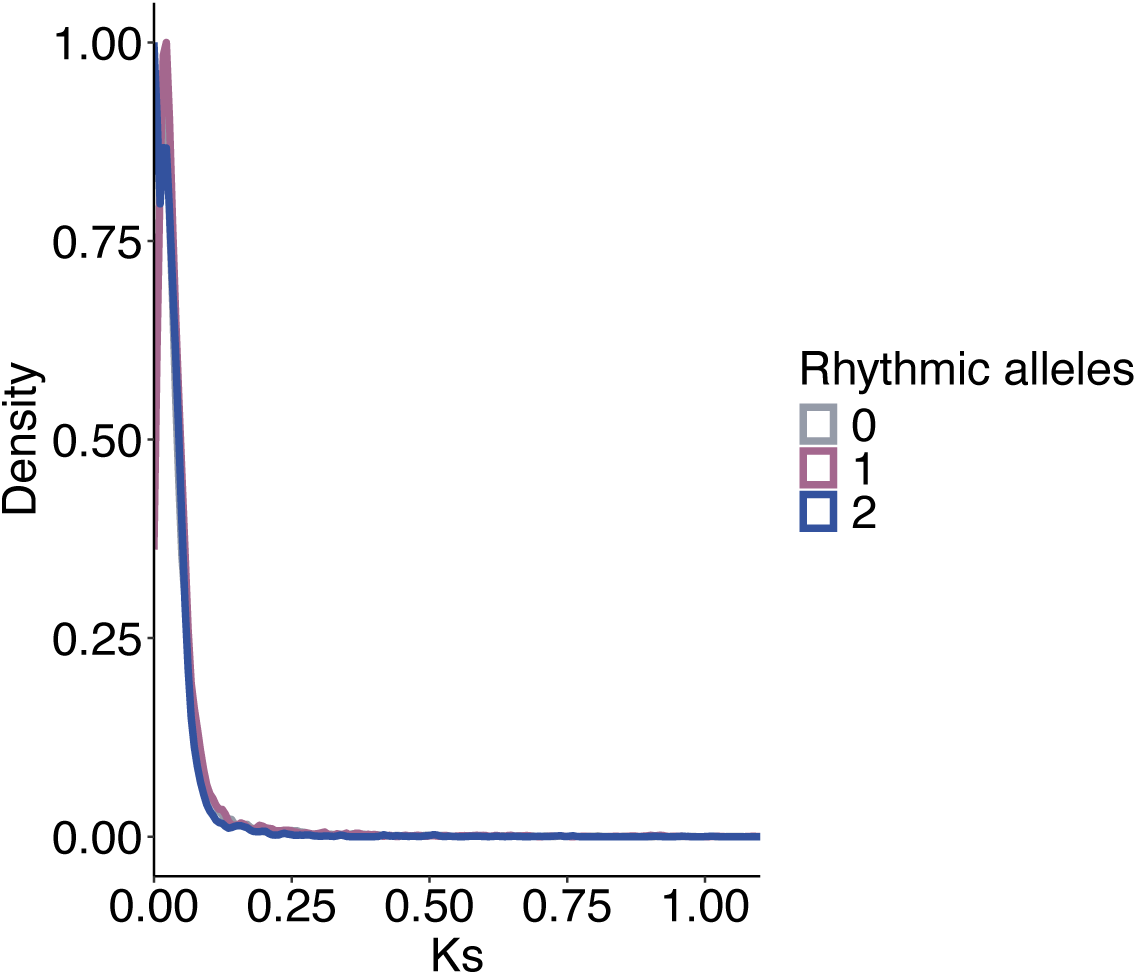
Distribution of Ks of syntenic allelic pairs of *Solanum tuberosum* cv. Atlantic, with either non, 1 or 2 rhythmic alleles as determined by JTK using the short day diel dataset.

**Supplemental Figure 3.**
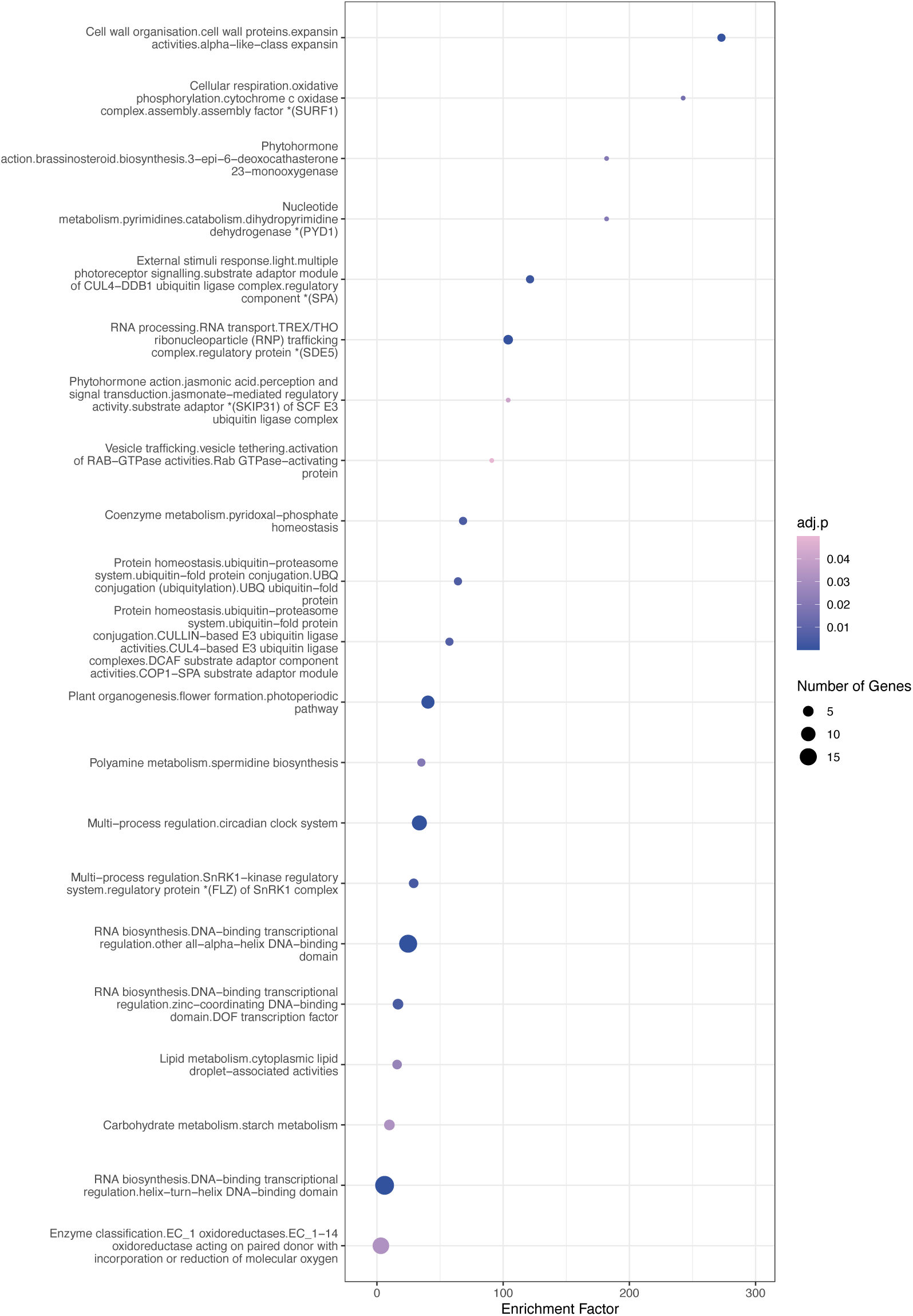
Functional enrichment of rhythmic transcripts that had a delayed phase in tubers with respect to leaves. Transcripts with at least 2 h delayed phase were compared to all tuber expressed transcripts.

**Supplemental Figure 4.**
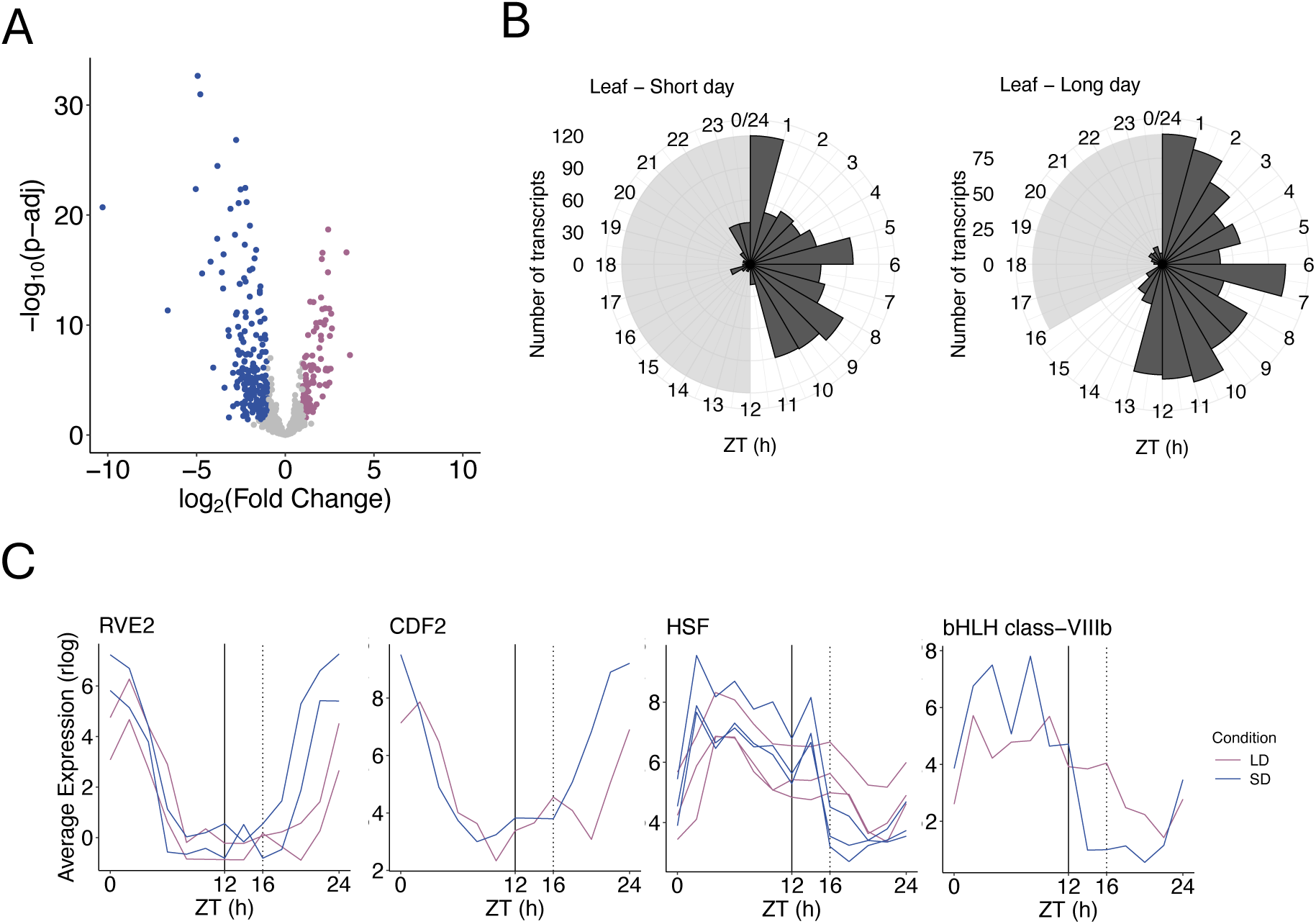
Expression characteristics of transcripts with larger amplitudes under short days. **A** Volcano plot of transcripts with larger amplitudes under SD than LD (DAmp>0.5). Blue indicates upregulated expression under SD and pink under LD. **B.** Phase of expression of genes with amplitudes higher in SD than LD (DAmp>0.5). **C.** Average leaf expression of transcriptional regulators with larger amplitudes under SD than LD (DAmp>1) (average n = 3 ).

**Supplemental Figure 5.**
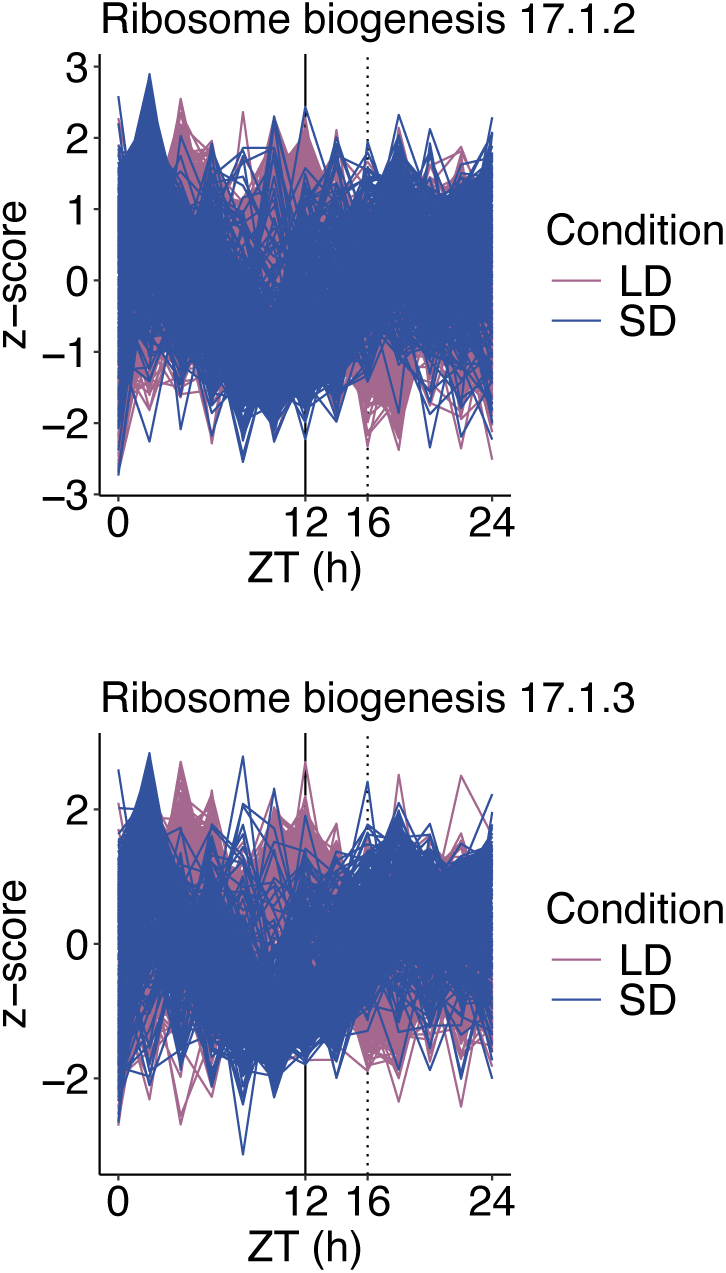
Expression patterns of all genes in the Mercator4 bin categories related to Protein biosynthesis.ribosome biogenesis.large ribosomal subunit (LSU) 17.1.2 and Protein biosynthesis.ribosome biogenesis.small ribosomal subunit (SSU) 17.1.3.

**Supplemental Figure 6.**
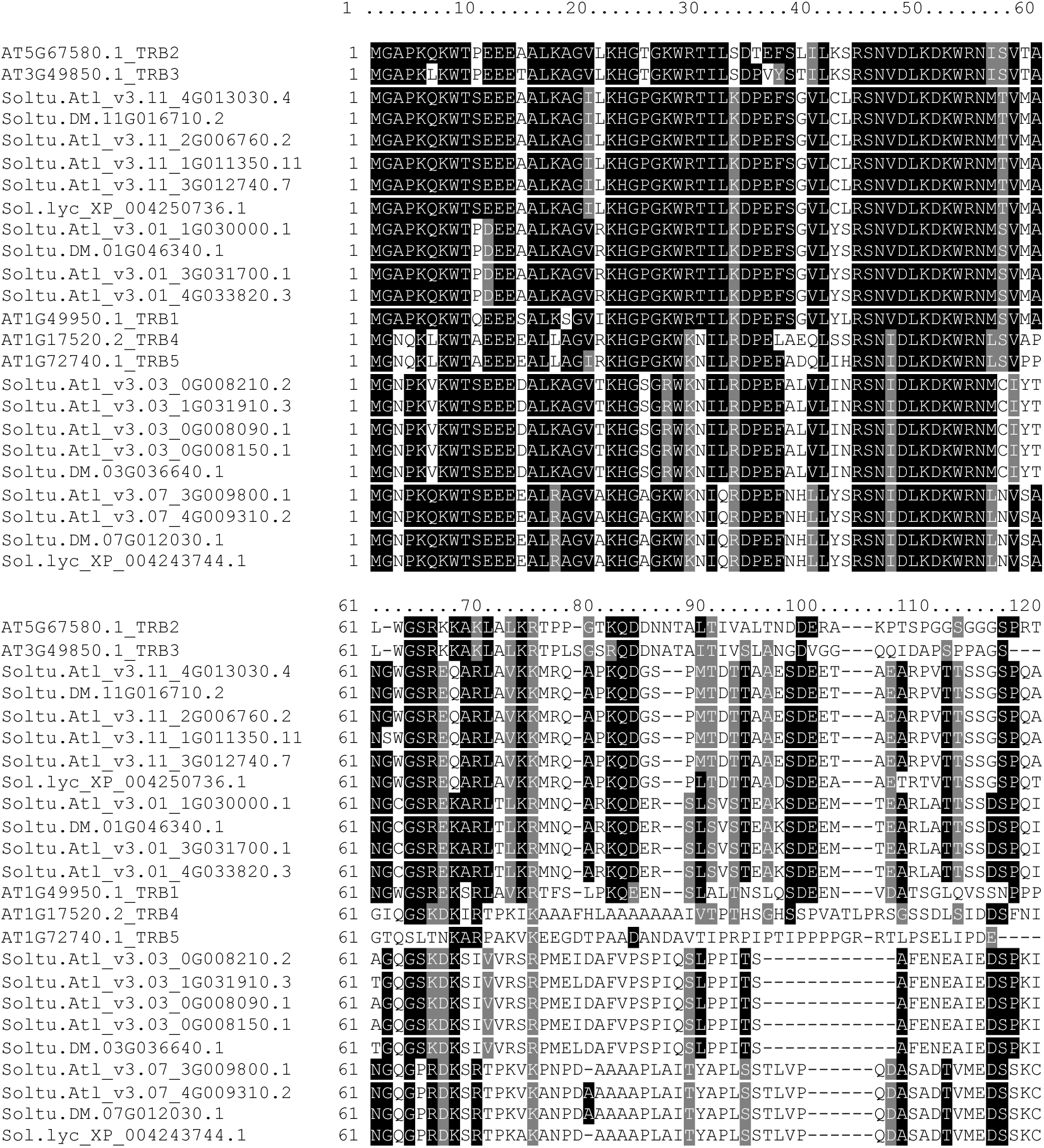
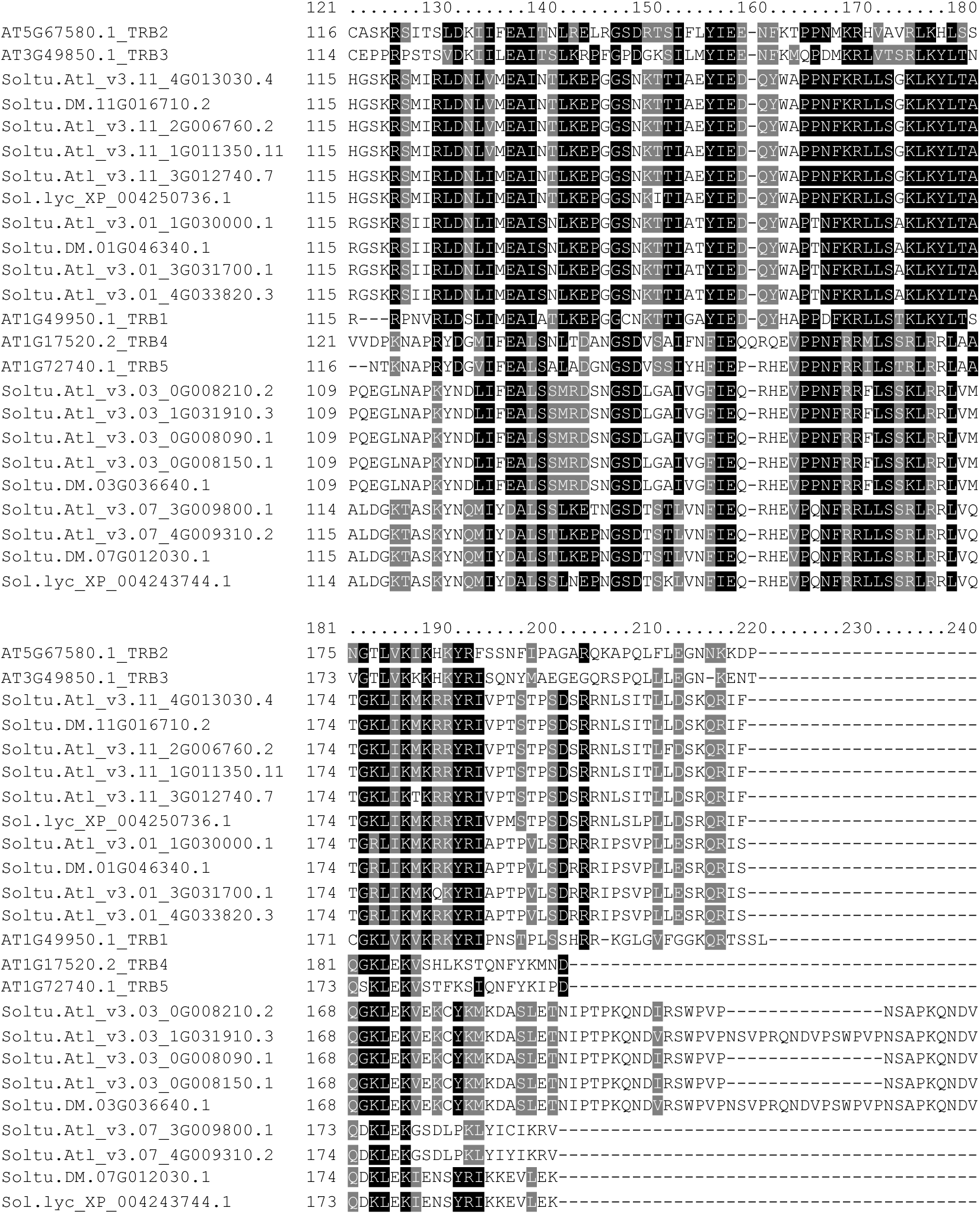
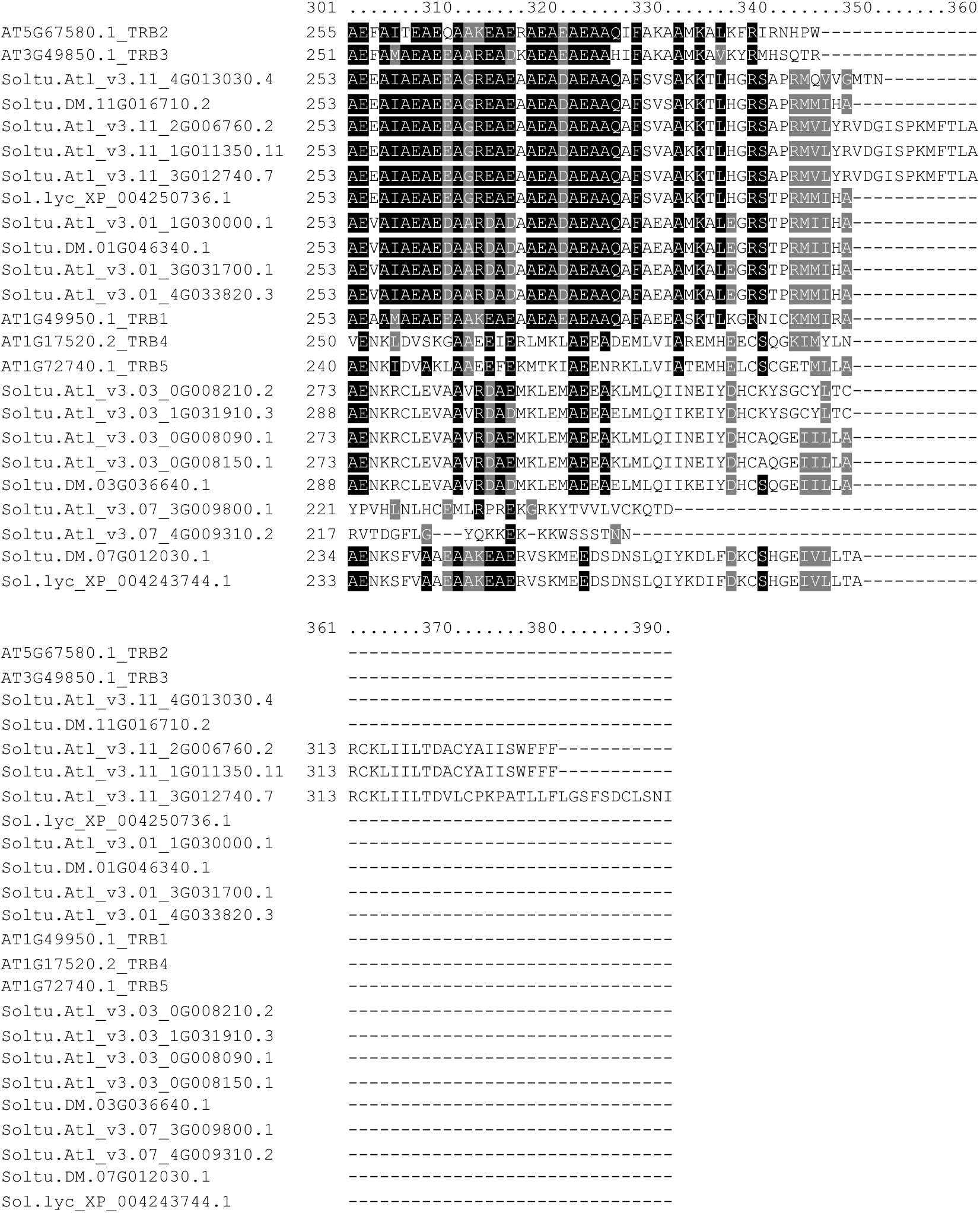
TRB proteins in Arabidopsis and potato. Peptide sequences were aligned using clustalW. Shading was implemented using BoxShade implemented using https://junli.netlify.app/apps/boxshade/.

**Supplemental Figure 7.**
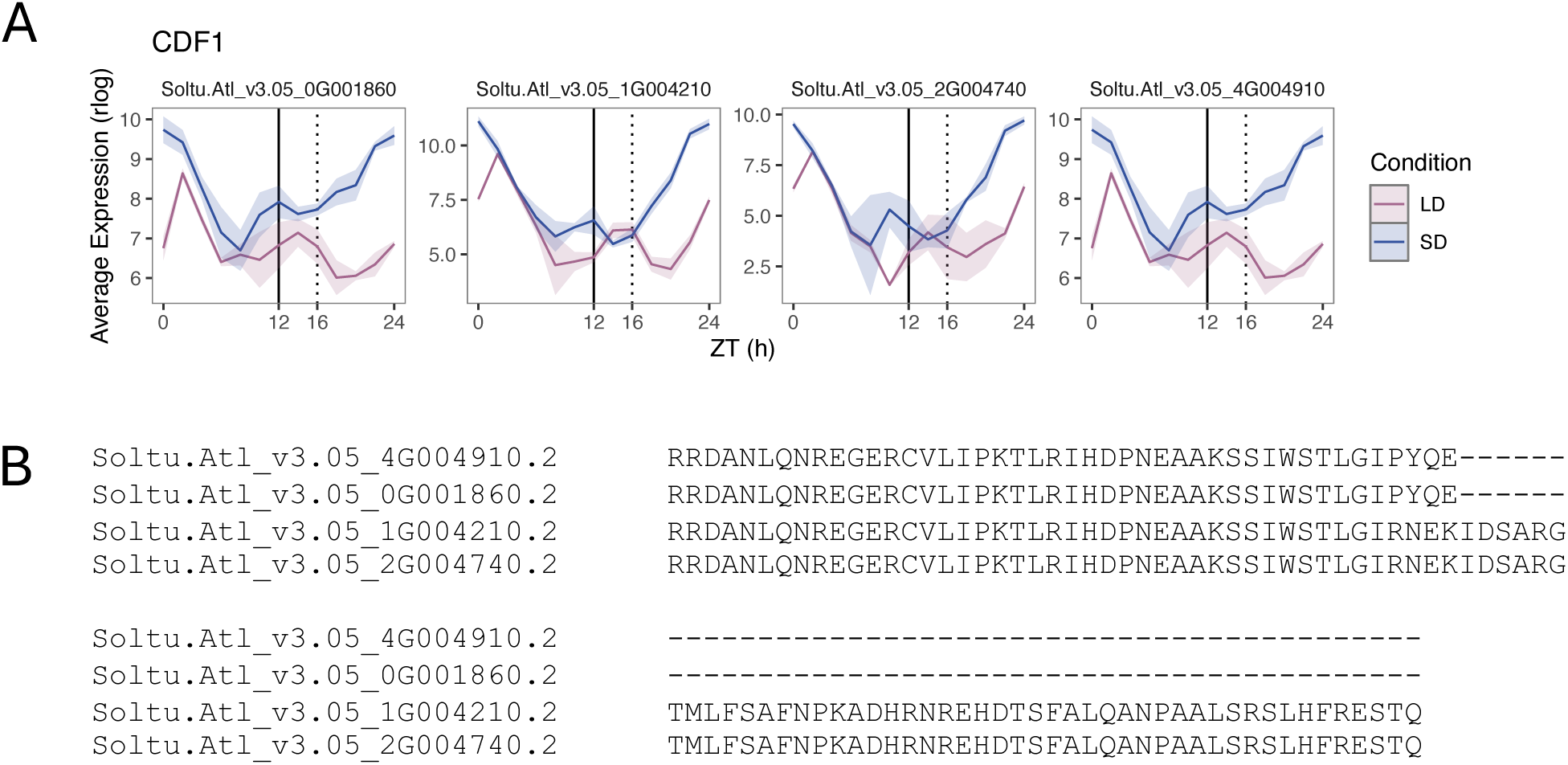
CDF1 alleles in Atlantic. A. Expression patterns *CDF1* alleles in short and long day conditions. Data represent the average ± standard deviation (n = 3 ). B. Alignment of the C-terminus end of CDF1 alleles.

**Supplemental Figure 8.**
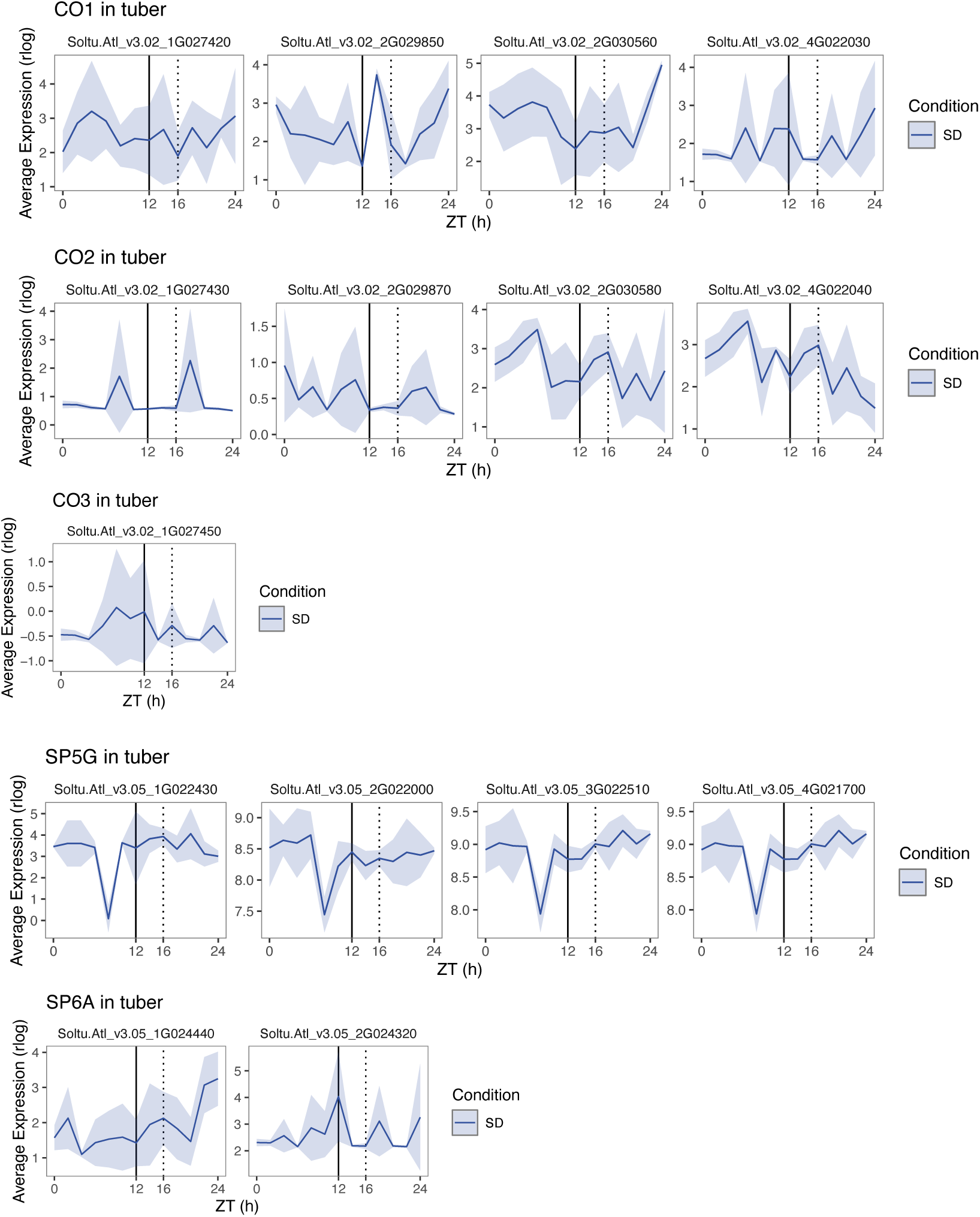
Expression patterns in tubers under short photoperiods. Data represents mean expression (average ± standard deviation, n = 3 ). SD, short days; LD, long days. *CO* genes include *CO1b* (Soltu.Atl_v3.02_2G030560), *CO2b* (Soltu.Atl_v3.02_2G030580), *CO3b* (Soltu.Atl_v3.02_2G030600).

**Supplemental Figure 9.**
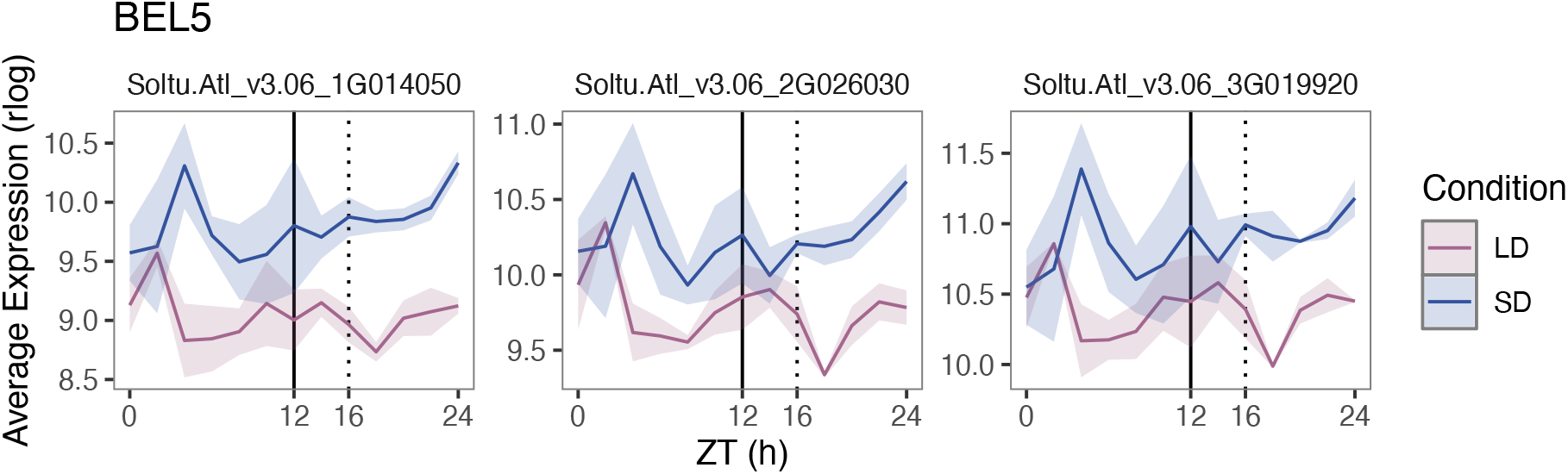
Expression patterns *BEL5* alleles in short and long day conditions. Data represent the average ± standard deviation (n = 3 ).

## REFERENCES

1. Salman-Minkov A, Sabath N, Mayrose I. Whole-genome duplication as a key factor in crop domestication. Nat Plants. 2016;2:16115.

2. Muller NA, Zhang L, Koornneef M, Jimenez-Gomez JM. Mutations in EID1 and LNK2 caused light-conditional clock deceleration during tomato domestication. Proc Natl Acad Sci U S A. 2018;115:7135–40.

3. Steed G, Ramirez DC, Hannah MA, Webb AAR. Chronoculture, harnessing the circadian clock to improve crop yield and sustainability. Science. 2021;372:eabc9141.

4. McClung CR. Circadian Clock Components Offer Targets for Crop Domestication and Improvement. Genes (Basel). 2021;12:374.

5. Hardigan MA, Laimbeer FPE, Newton L, Crisovan E, Hamilton JP, Vaillancourt B, et al. Genome diversity of tuber-bearing Solanum uncovers complex evolutionary history and targets of domestication in the cultivated potato. Proc Natl Acad Sci U S A. 2017;114:E9999–10008.

6. Lysak MA, Koch MA, Pecinka A, Schubert I. Chromosome triplication found across the tribe Brassiceae. Genome Res. 2005;15:516–25.

7. Beilstein MA, Nagalingum NS, Clements MD, Manchester SR, Mathews S. Dated molecular phylogenies indicate a Miocene origin for Arabidopsis thaliana. Proc Natl Acad Sci U S A. 2010;107:18724–8.

8. Yang YW, Lai KN, Tai PY, Li WH. Rates of nucleotide substitution in angiosperm mitochondrial DNA sequences and dates of divergence between Brassica and other angiosperm lineages. J Mol Evol. 1999;48:597–604.

9. Levy AA, Feldman M. Evolution and origin of bread wheat. The Plant Cell. 2022;34:2549–67.

10. Greenham K, Sartor RC, Zorich S, Lou P, Mockler TC, McClung CR. Expansion of the circadian transcriptome in Brassica rapa and genome-wide diversification of paralog expression patterns. Elife. 2020;9:e58993.

11. Rees H, Rusholme-Pilcher R, Bailey P, Colmer J, White B, Reynolds C, et al. Circadian regulation of the transcriptome in a complex polyploid crop. PLoS Biol. 2022;20:e3001802.

12. Jansky SH, Spooner DM. The Evolution of Potato Breeding. In: Goldman I, editor. Plant Breeding Reviews. 1st edition. Wiley; 2018. p. 169–214.

13. Gutaker RM, Weiß CL, Ellis D, Anglin NL, Knapp S, Luis Fernández-Alonso J, et al. The origins and adaptation of European potatoes reconstructed from historical genomes. Nature Ecology & Evolution. 2019;3:1093–101.

14. Zhang C, Wang P, Tang D, Yang Z, Lu F, Qi J, et al. The genetic basis of inbreeding depression in potato. Nat Genet. 2019;51:374–8.

15. Hijmans RJ. Global distribution of the potato crop. Am J Pot Res. 2001;78:403–12.

16. Abelenda JA, Cruz-Oro E, Franco-Zorrilla JM, Prat S. Potato StCONSTANS-like1 Suppresses Storage Organ Formation by Directly Activating the FT-like StSP5G Repressor. Curr Biol. 2016;26:872–81.

17. Navarro C, Abelenda JA, Cruz-Oro E, Cuellar CA, Tamaki S, Silva J, et al. Control of flowering and storage organ formation in potato by FLOWERING LOCUS T. Nature. 2011;478:119–22.

18. Kloosterman B, Abelenda JA, Gomez Mdel M, Oortwijn M, de Boer JM, Kowitwanich K, et al. Naturally occurring allele diversity allows potato cultivation in northern latitudes. Nature. 2013;495:246–50.

19. Hoopes GM, Zarka D, Feke A, Acheson K, Hamilton JP, Douches D, et al. Keeping time in the dark: Potato diel and circadian rhythmic gene expression reveals tissue-specific circadian clocks. Plant Direct. 2022;6:e425.

20. Hoopes G, Meng X, Hamilton JP, Achakkagari SR, de Alves Freitas Guesdes F, Bolger ME, et al. Phased, chromosome-scale genome assemblies of tetraploid potato reveal a complex genome, transcriptome, and predicted proteome landscape underpinning genetic diversity. Mol Plant. 2022;15:520–36.

21. Hughes ME, Hogenesch JB, Kornacker K. JTK_CYCLE: an efficient nonparametric algorithm for detecting rhythmic components in genome-scale data sets. J Biol Rhythms. 2010;25:372–80.

22. Kryuchkova-Mostacci N, Robinson-Rechavi M. A benchmark of gene expression tissue-specificity metrics. Briefings in Bioinformatics. 2017;18:205–14.

23. Sun H, Jiao W-B, Krause K, Campoy JA, Goel M, Folz-Donahue K, et al. Chromosome-scale and haplotype-resolved genome assembly of a tetraploid potato cultivar. Nat Genet. 2022;54:342–8.

24. Bao Z, Li C, Li G, Wang P, Peng Z, Cheng L, et al. Genome architecture and tetrasomic inheritance of autotetraploid potato. Molecular Plant. 2022;15:1211–26.

25. Wang F, Xia Z, Zou M, Zhao L, Jiang S, Zhou Y, et al. The autotetraploid potato genome provides insights into highly heterozygous species. Plant Biotechnol J. 2022;20:1996–2005.

26. Endo M. Tissue-specific circadian clocks in plants. Curr Opin Plant Biol. 2016;29:44–9.

27. James AB, Monreal JA, Nimmo GA, Kelly CL, Herzyk P, Jenkins GI, et al. The circadian clock in Arabidopsis roots is a simplified slave version of the clock in shoots. Science. 2008;322:1832–5.

28. Takahashi N, Hirata Y, Aihara K, Mas P. A hierarchical multi-oscillator network orchestrates the Arabidopsis circadian system. Cell. 2015;163:148–59.

29. Bordage S, Sullivan S, Laird J, Millar AJ, Nimmo HG. Organ specificity in the plant circadian system is explained by different light inputs to the shoot and root clocks. New Phytol. 2016;212:136–49.

30. Michael TP, Mockler TC, Breton G, McEntee C, Byer A, Trout JD, et al. Network discovery pipeline elucidates conserved time-of-day-specific cis-regulatory modules. PLoS Genet. 2008;4:e14.

31. Seaton DD, Graf A, Baerenfaller K, Stitt M, Millar AJ, Gruissem W. Photoperiodic control of the Arabidopsis proteome reveals a translational coincidence mechanism. Mol Syst Biol. 2018;14:e7962.

32. Wang Q, Liu W, Leung CC, Tarté DA, Gendron JM. Plants distinguish different photoperiods to independently control seasonal flowering and growth. Science. 2024;383:eadg9196.

33. Schrumpfová PP, Vychodilová I, Hapala J, Schořová Š, Dvořáček V, Fajkus J. Telomere binding protein TRB1 is associated with promoters of translation machinery genes in vivo. Plant Mol Biol. 2016;90:189–206.

34. Abelenda JA, Navarro C, Prat S. Flowering and tuberization: a tale of two nightshades. Trends Plant Sci. 2014;19:115–22.

35. Morris WL, Hancock RD, Ducreux LJ, Morris JA, Usman M, Verrall SR, et al. Day length dependent restructuring of the leaf transcriptome and metabolome in potato genotypes with contrasting tuberization phenotypes. Plant Cell Environ. 2014;37:1351–63.

36. Ferrari C, Proost S, Janowski M, Becker J, Nikoloski Z, Bhattacharya D, et al. Kingdom-wide comparison reveals the evolution of diurnal gene expression in Archaeplastida. Nature Communications. 2019;10.

37. Panchy N, Wu G, Newton L, Tsai CH, Chen J, Benning C, et al. Prevalence, evolution, and cis-regulation of diel transcription in Chlamydomonas reinhardtii. G3 (Bethesda). 2014;4:2461–71.

38. Wang G-Z, Hickey SL, Shi L, Huang H-C, Nakashe P, Koike N, et al. Cycling Transcriptional Networks Optimize Energy Utilization on a Genome Scale. Cell Rep. 2015;13:1868–80.

39. Ramos A, Perez-Solis E, Ibanez C, Casado R, Collada C, Gomez L, et al. Winter disruption of the circadian clock in chestnut. Proc Natl Acad Sci U S A. 2005;102:7037–42.

40. Nose M, Watanabe A. Clock genes and diurnal transcriptome dynamics in summer and winter in the gymnosperm Japanese cedar (Cryptomeria japonica (L.f.) D.Don). BMC Plant Biol. 2014;14:308.

41. Jung S, Kim H, Lee J, Kang MH, Kim J, Kim JK, et al. The genetically programmed rhythmic alteration of diurnal gene expression in the aged Arabidopsis leaves. Front Plant Sci. 2024;15:1481682.

42. Li Z, Bonaldi K, Uribe F, Pruneda-Paz JL. A Localized Pseudomonas syringae Infection Triggers Systemic Clock Responses in Arabidopsis. Curr Biol. 2018;28:630–639.e4.

43. Xu Y, Koroma AA, Weise SE, Fu X, Sharkey TD, Shachar-Hill Y. Daylength variation affects growth, photosynthesis, leaf metabolism, partitioning, and metabolic fluxes. Plant Physiol. 2023;194:475–90.

44. Missra A, Ernest B, Lohoff T, Jia Q, Satterlee J, Ke K, et al. The Circadian Clock Modulates Global Daily Cycles of mRNA Ribosome Loading. The Plant Cell. 2015;27:2582–99.

45. Amiard S, Feit L, Vanrobays E, Simon L, Le Goff S, Loizeau L, et al. The TELOMERE REPEAT BINDING proteins TRB4 and TRB5 function as transcriptional activators of PRC2- controlled genes to regulate plant development. Plant Communications. 2024;5:100890.

46. Griebel T, Zeier J. Light Regulation and Daytime Dependency of Inducible Plant Defenses in Arabidopsis: Phytochrome Signaling Controls Systemic Acquired Resistance Rather Than Local Defense. Plant Physiol. 2008;147:790–801.

47. Bhardwaj V, Meier S, Petersen LN, Ingle RA, Roden LC. Defence Responses of Arabidopsis thaliana to Infection by Pseudomonas syringae Are Regulated by the Circadian Clock. PLoS One. 2011;6:e26968.

48. Cortleven A, Roeber VM, Frank M, Bertels J, Lortzing V, Beemster GTS, et al. Photoperiod Stress in Arabidopsis thaliana Induces a Transcriptional Response Resembling That of Pathogen Infection. Front Plant Sci. 2022;13:838284.

49. Ingle RA, Stoker C, Stone W, Adams N, Smith R, Grant M, et al. Jasmonate signalling drives time-of-day differences in susceptibility of Arabidopsis to the fungal pathogen Botrytis cinerea. Plant J. 2015;84:937–48.

50. Zhang C, Wang D, Yang C, Kong N, Shi Z, Zhao P, et al. Genome-wide identification of the potato WRKY transcription factor family. PLOS ONE. 2017;12:e0181573.

51. Villano C, Esposito S, D’Amelia V, Garramone R, Alioto D, Zoina A, et al. WRKY genes family study reveals tissue-specific and stress-responsive TFs in wild potato species. Sci Rep. 2020;10:7196.

52. Plantenga FDM, Heuvelink E, Rienstra JA, Visser RGF, Bachem CWB, Marcelis LFM. Coincidence of potato CONSTANS (StCOL1) expression and light cannot explain night-break repression of tuberization. Physiol Plant. 2019;167:250–63.

53. González-Schain ND, Díaz-Mendoza M, Zurczak M, Suárez-López P. Potato CONSTANS is involved in photoperiodic tuberization in a graft-transmissible manner. Plant J. 2012;70:678–90.

54. Sharma P, Lin T, Hannapel DJ. Targets of the StBEL5 Transcription Factor Include the FT Ortholog StSP6A. Plant Physiol. 2016;170:310–24.

55. Shaikh MA, Ramírez-Gonzales L, Franco-Zorrilla JM, Steiner E, Oortwijn M, Bachem CWB, et al. StCDF1: A “jack of all trades” clock output with a central role in regulating potato nitrate reduction activity. New Phytol. 2025;245:282–98.

56. Wan CY, Wilkins TA. A modified hot borate method significantly enhances the yield of high-quality RNA from cotton (Gossypium hirsutum L.). Anal Biochem. 1994;223:7–12.

57. Wood DE, Lu J, Langmead B. Improved metagenomic analysis with Kraken 2. Genome Biology. 2019;20:257.

58. Martin M. Cutadapt removes adapter sequences from high-throughput sequencing reads. EMBnet j. 2011;17:10.

59. Ewels P, Magnusson M, Lundin S, Käller M. MultiQC: summarize analysis results for multiple tools and samples in a single report. Bioinformatics. 2016;32:3047–8.

60. Bray NL, Pimentel H, Melsted P, Pachter L. Near-optimal probabilistic RNA-seq quantification. Nat Biotechnol. 2016;34:525–7.

61. Love MI, Huber W, Anders S. Moderated estimation of fold change and dispersion for RNA-seq data with DESeq2. Genome Biology. 2014;15:550.

62. Wu G, Anafi RC, Hughes ME, Kornacker K, Hogenesch JB. MetaCycle: an integrated R package to evaluate periodicity in large scale data. Bioinformatics. 2016;32:3351–3.

63. Virtanen P, Gommers R, Oliphant TE, Haberland M, Reddy T, Cournapeau D, et al. SciPy 1.0: fundamental algorithms for scientific computing in Python. Nat Methods. 2020;17:261–72.

64. McKinney W. Data Structures for Statistical Computing in Python. In: Walt S van der, Millman J, editors. Proceedings of the 9th Python in Science Conference. 2010. p. 56–61.

65. Lovell JT, Sreedasyam A, Schranz ME, Wilson M, Carlson JW, Harkess A, et al. GENESPACE tracks regions of interest and gene copy number variation across multiple genomes. eLife. 2022;11:e78526.

66. Hamilton JP, Brose J, Buell CR. SpudDB: a database for accessing potato genomic data. Genetics. 2025;229:iyae205.

67. Albert VA, Krabbenhoft TJ. Navigating the CoGe Online Software Suite for Polyploidy Research. In: Van de Peer Y, editor. Polyploidy: Methods and Protocols. New York, NY: Springer US; 2023. p. 19–45.

68. Bolger M, Schwacke R, Usadel B. MapMan Visualization of RNA-Seq Data Using Mercator4 Functional Annotations. Methods Mol Biol. 2021;2354:195–212.

69. Heinz S, Benner C, Spann N, Bertolino E, Lin YC, Laslo P, et al. Simple Combinations of Lineage-Determining Transcription Factors Prime cis-Regulatory Elements Required for Macrophage and B Cell Identities. Molecular Cell. 2010;38:576–89.

70. Jin J, Tian F, Yang D-C, Meng Y-Q, Kong L, Luo J, et al. PlantTFDB 4.0: toward a central hub for transcription factors and regulatory interactions in plants. Nucleic Acids Research. 2017;45:D1040–5.

